# TACE reprograms RANKL-mediated differentiation of macrophages by activating the non-canonical pathway of IRF3

**DOI:** 10.64898/2026.02.11.705073

**Authors:** Se Hwan Mun, Brian Oh, Shunichi Yokota, Akio Umemoto, Andrew Suh, Kyuho Kang, Woojung Kim, David Oliver, Tania Pannellini, Geunho Kwon, Young Yang, Liang Deng, Kyung-Hyun Park-Min

## Abstract

Inflammation is associated with an influx of inflammatory macrophages and increased activation and differentiation of osteoclasts. Receptor activator of NF-kB ligand (RANKL) is a key driver for osteoclast differentiation. However, the pathogenic mechanisms augmenting RANKL-induced osteoclast differentiation in inflammatory conditions are not fully elucidated. Here, we show that TNF-α converting enzyme (TACE) plays a critical role in pathological bone erosion and enhances osteoclast differentiation in inflammatory conditions. Myeloid cell-specific TACE deletion in a murine arthritis model attenuates joint inflammation and bone destruction. TACE deficiency suppresses distal RANKL signaling in macrophages by increasing IRF3 activation, while enhancing proximal RANKL signaling, leading to the suppression of osteoclast differentiation. Mechanistically, IRF3 activation limits macrophage reprogramming by suppressing NFATc1 and HB-EGF in response to RANKL through a non-canonical pathway. HB-EGF, a TACE substrate, activates EGFR signaling and promotes osteoclastogenesis by inhibiting IRF3 activation. TACE regulates the reciprocal inhibition of the IRF3-HBEGF axis. Our study highlights a role for TACE as a rheostat, balancing both pro- and anti-osteoclastogenic signals.

## Introduction

Inflammatory osteolysis is associated with the activation of osteoclasts and found in many inflammatory diseases, including rheumatoid arthritis (RA). RA is a chronic inflammatory autoimmune disease^1, 2^. Progressive bone erosion and joint destruction are key clinical features of RA that lead to increased risk of fractures and severe disabilities ^3^. Arthritic bone erosion is driven by imbalanced bone remodeling and excessive osteoclast-mediated bone resorption; osteoclasts are the sole bone-resorbing cells and a key driver of steady-state bone remodeling and inflammatory bone loss^4, 5, 6^. Alleviating bone erosion holds importance for treating RA. Several pathological mechanisms of bone erosion in RA have been identified^7^. However, the underlying mechanism of arthritic bone erosion remains insufficiently characterized.

Osteoclasts are derived from monocyte/macrophage lineage cells^8, 9^. RA synovial fluid macrophages consist of multiple populations that differentiate into osteoclasts ^10, 11, 12^. Altered gene and protein expressions in RA synovial CD14+ cells have been proposed to activate osteoclasts ^13^. Macrophage colony-stimulating factor (M-CSF) and receptor activator of NF-κB ligand (RANKL) are key factors for osteoclastogenesis^14^. RANKL activates multiple downstream signaling pathways, including NF-kB and MAPK signaling pathways, and subsequently induces the expression of NFATc1, a master regulator of osteoclastogenesis ^15, 16, 17^. The contribution of inflammatory cytokines to promoting pathogenic osteoclasts has been highlighted ^18, 19, 20^. Although the mechanisms responsible for the aberrant activation of joint erosion by osteoclasts are complex, the treatment of RA patients with denosumab, an anti-RANKL therapy, shows increased bone mineral density and diminished progression of joint destruction with a limited impact on articular inflammation^21, 22^. However, how RANKL signaling is amplified to promote osteoclastogenesis in inflammatory diseases is poorly understood.

Under homeostatic conditions, osteoclasts are controlled by the balance of positive and negative signals^6^. While the positive signals for osteoclastogenesis have been extensively studied^6^, a few negative feedback mechanisms, including the type I and type II interferons (IFNs), have been identified so far. IFN signaling protects mice from bone loss and restricts osteoclastogenesis through activation of negative feedback signaling^23^. Although the negative effect of IFNβ on osteoclast formation is evident^24^, the underlying mechanism of type I IFN-mediated negative regulation has not been fully elucidated. IRF3 (Interferon Regulatory Factor 3) is a transcription factor that plays a crucial role in the production of type I interferons in response to viral and bacterial infections ^25^. Upon activation, IRF3 undergoes phosphorylation and forms homo- or heterodimers that translocate to the nucleus, where they bind to the enhancer regions of the IFNβ gene^26^ and further induce interferon-regulated genes (ISGs). IRF3 can be activated in macrophages by various signaling pathways, such as the STING pathway^27, 28, 29^. However, the direct role of IRF3 in osteoclast differentiation has not been studied. Our study determined the novel function of the IRF3 non-canonical pathway in osteoclast formation and bone remodeling.

TNF-alpha converting enzyme (TACE, also known as ADAM17) is a transmembrane metalloprotease that triggers ectodomain shedding of cell surface proteins such as TNF-α, Notch, and the heparin-binding epidermal growth factor (EGF)-like growth factor (HB-EGF) ^30, 31,32^. By regulating the shedding of receptors or ligands, TACE is involved in diverse cellular functions, including rapid protein down-regulation, response to cytokines, secretion of cytokines, immune responses, and cancer development^30, 31, 32^. TNF-α is a major substrate of TACE and is involved in synovial inflammation of inflammatory arthritis^33^. TACE is required for osteoclast migration during embryonic development^34^. TACE deletion in hematopoietic stem cells negatively regulates the size of osteoclasts without affecting osteoclast differentiation^35^. The role of TACE in osteoclasts is conflicting and remains unclear. TACE expression in synovial macrophages of RA patients is significantly increased. Here, we examined the role of TACE in osteoclast differentiation and bone erosion in inflammatory arthritis using myeloid-cell-specific TACE-deficient TNF transgenic mice. Myeloid-specific TACE deletion attenuates arthritic bone erosion and osteoclast formation in mice with inflammatory arthritis. TACE positively regulates both human and mouse osteoclast differentiation. Consistent with its sheddase role, TACE-deficiency increases the proximal RANKL signaling. However, TACE deficiency inhibits distal RANKL signaling by activating IRF3, which is mediated by the non-canonical pathway of IRF3 and occurs independently of IFNs. While IRF3 is involved in an IFN-dependent expression of ISGs by RANKL stimulation, IRF3 also suppresses positive regulators of osteoclasts, such as NFATc1, epidermal growth factor receptor (EGFR), and HB-EGF, in an IFN-independent manner. Supplementing HB-EGF promotes osteoclastogenesis and suppresses RANKL-induced IRF3 activation. TACE and HB-EGF expression are elevated in macrophages of synovial fluids and synovium from RA patients, indicating a clinical relevance of the TACE-IRF3-EGFR axis in RANKL-induced osteoclastogenesis. Our data reveal an unrecognized role of metalloproteases in promoting RANKL-induced osteoclast differentiation during both homeostatic and pathological bone resorption.

## Results

### TACE deficiency inhibits RANKL-mediated osteoclastogenesis

We aimed to identify a new pathway to promote RANKL-mediated osteoclast formation in inflammatory bone diseases, such as rheumatoid arthritis (RA). As osteoclasts are derived from synovial fluid (SF) CD14+ macrophages of patients with RA ^36^, the expression of TACE in SF CD14+ macrophages was assessed. TACE expression was significantly upregulated in SF CD14+ macrophages from RA patients (Supplementary Fig. 1a). To identify the mechanism responsible for increased TACE expression in SF macrophages, we used a well-established system for human macrophages; human peripheral blood CD14+ monocytes were cultured with M-CSF. We treated human CD14+ cells with TNF-α or RANKL, which are abundant in RA synovial fluids and key inducers of osteoclast formation^37, 38^. Both TNF-α and RANKL increased TACE expression in CD14+ cells (Supplementary Fig. 1b, c), indicating that the inflammatory conditions prime macrophages to induce TACE expression. To determine the effect of TACE on osteoclast differentiation, we knocked down TACE in human CD14+ cells using small interfering RNAs (siRNAs). We first verified the efficiency of knockdown (KD) of TACE mRNA and protein in human CD14+ cells (Fig. 1a, b). Human CD14+ monocytes were cultured with M-CSF and RANKL for four days (Supplementary Fig. 1d), and *in vitro* osteoclast differentiation was evaluated by counting multinucleated tartrate-resistant acid phosphatase (TRAP)-positive multinuclear cells. TACE knockdown effectively suppressed RANKL-mediated osteoclast formation compared to control cells (Fig. 1c). Accordingly, RANKL-induced expression of osteoclast marker genes, including NFATC1 (encoding NFATc1), CTSK (encoding cathepsin K), and ITGB3 (encoding integrin β3), was suppressed in TACE KD cells relative to control cells (Fig. 1d, e). To corroborate our findings, TACE was overexpressed by transduction with adenoviral particles encoding TACE or GFP. Adenoviral transduction induced TACE expression (Fig. 1f). Ectopic expression of TACE enhanced osteoclast differentiation (Fig. 1g) and the expression of NFATc1, an osteoclast marker gene (Fig. 1h). Together, these results indicate that TACE positively regulates RANKL-mediated osteoclast differentiation.

**Fig. 1.**
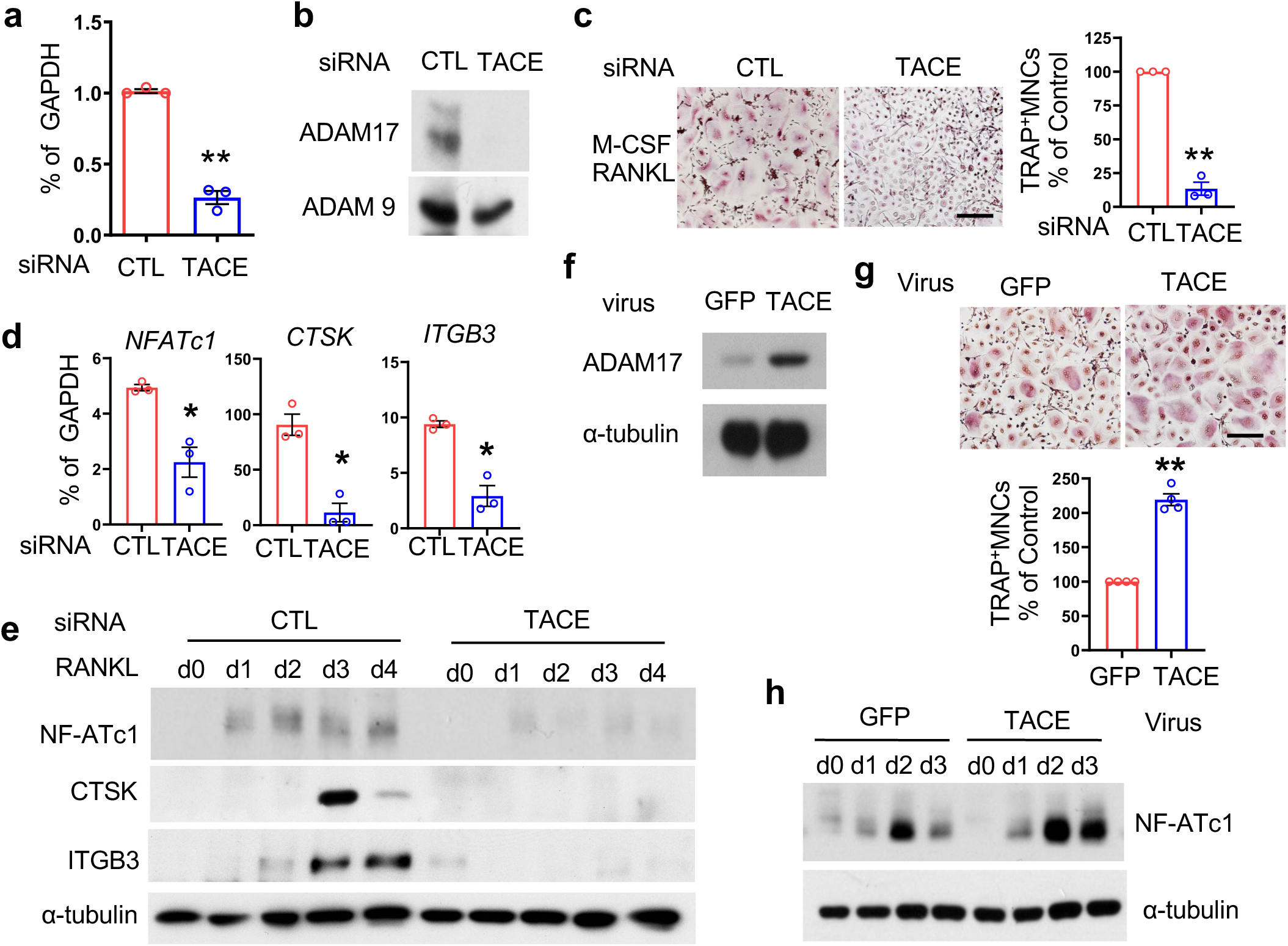
TACE deficiency inhibits human osteoclast formation. (**a**-**d**) Human CD14⁺ cells were nucleofected with either control (CTL) or TACE-targeting siRNAs and cultured with M-CSF. (**a**,**b**) Knockdown (KD) efficiency. (**a**) TACE mRNA levels were measured by qPCR and normalized to GAPDH. (**b**) Immunoblot analysis of KD cells using an anti-TACE antibody. (**c**, **d**) CTL and TACE KD CD14⁺ cells were cultured with M-CSF (20 ng/ml) and RANKL (40 ng/ml) for 5 days.(**c**) Osteoclastogenesis assay. Left: TRAP staining of representative images. Scale bar: 100 µm. Right: Quantification of TRAP-positive multinucleated cells (≥3 nuclei), normalized to CTL siRNA conditions. (**d**) RT-qPCR analysis of osteoclast-specific genes in CTL and TACE KD cells, normalized to GAPDH.(E) Immunoblot analysis using anti-NFATc1, Cathepsin K (CTSK), Integrin β3 (ITGB3), and α-tubulin antibodies.(**f**–**h**) Human CD14⁺ cells were transduced with either control-GFP or TACE-expressing adenovirus and cultured with M-CSF (20 ng/ml) and RANKL (40 ng/ml) for 5 days. (**f**) Osteoclastogenesis assay. Upper: TRAP staining of representative images. Scale bar: 100 µm. Bottom: Quantification of TRAP-positive multinucleated cells. (**h**) Immunoblot analysis using anti-NFATc1 and α-tubulin antibodies. All data are presented as mean ± SEM. *p < 0.05, **p < 0.01 by two-tailed, unpaired *t*-test (**a**, **c**, **d**, **g**). Data represent at least three independent experiments.

### TACE deficiency attenuates inflammatory bone destruction

We next assessed the impact of TACE on arthritic bone erosion. To account for the potential defect of endogenous TNF-α secretion by TACE-deficiency, we investigated the role of TACE in inflammatory bone destruction using a TNF transgene (tg)-induced arthritis model. We crossed myeloid-specific TACE-deficient mice with TNF transgenic mice to generate TNF transgenic mice with myeloid-specific TACE deficiency (called TACE^ΔM^ TNFtg) (Fig. 2a). TACE expression in bone marrow-derived macrophages (BMDMs) from TACE^ΔM^ TNFtg mice was significantly diminished, and both control TNF transgenic mice (named TACE^con^ tg) and TACE^ΔM^ TNFtg mice exhibited similar characteristics to control TNF transgenic mice (Supplementary Fig. 2a-e). We measured ankle swelling in TACE^con^ TNFtg and TACE^ΔM^ TNFtg mice and found a decrease in ankle swelling in TACE^ΔM^ TNFtg mice compared to TACE^con^ TNFtg mice (Fig. 2b). Human and mouse TNF-α levels in serum were measured. As expected, both TACE^con^ TNFtg and TACE^ΔM^ TNFtg mice exhibited high levels of serum human TNF-α, which were comparable between groups (Fig. 2c). However, serum mouse TNF-α in our system was undetectable (data not shown). Micro-CT analysis of ankle joints revealed that TACE^ΔM^ TNFtg mice were protected against bone erosion compared to TACE^con^ TNFtg mice (Fig. 2d). Consistently, histomorphometric analysis of the ankle in TACE^ΔM^ TNFtg mice showed a significant decrease in inflammation and bone erosion compared to the control mice (Fig. 2e). These results collectively demonstrate that TACE deficiency attenuated inflammatory bone destruction and ankle joint deformity in a TNFtg-induced arthritis model.

**Fig. 2.**
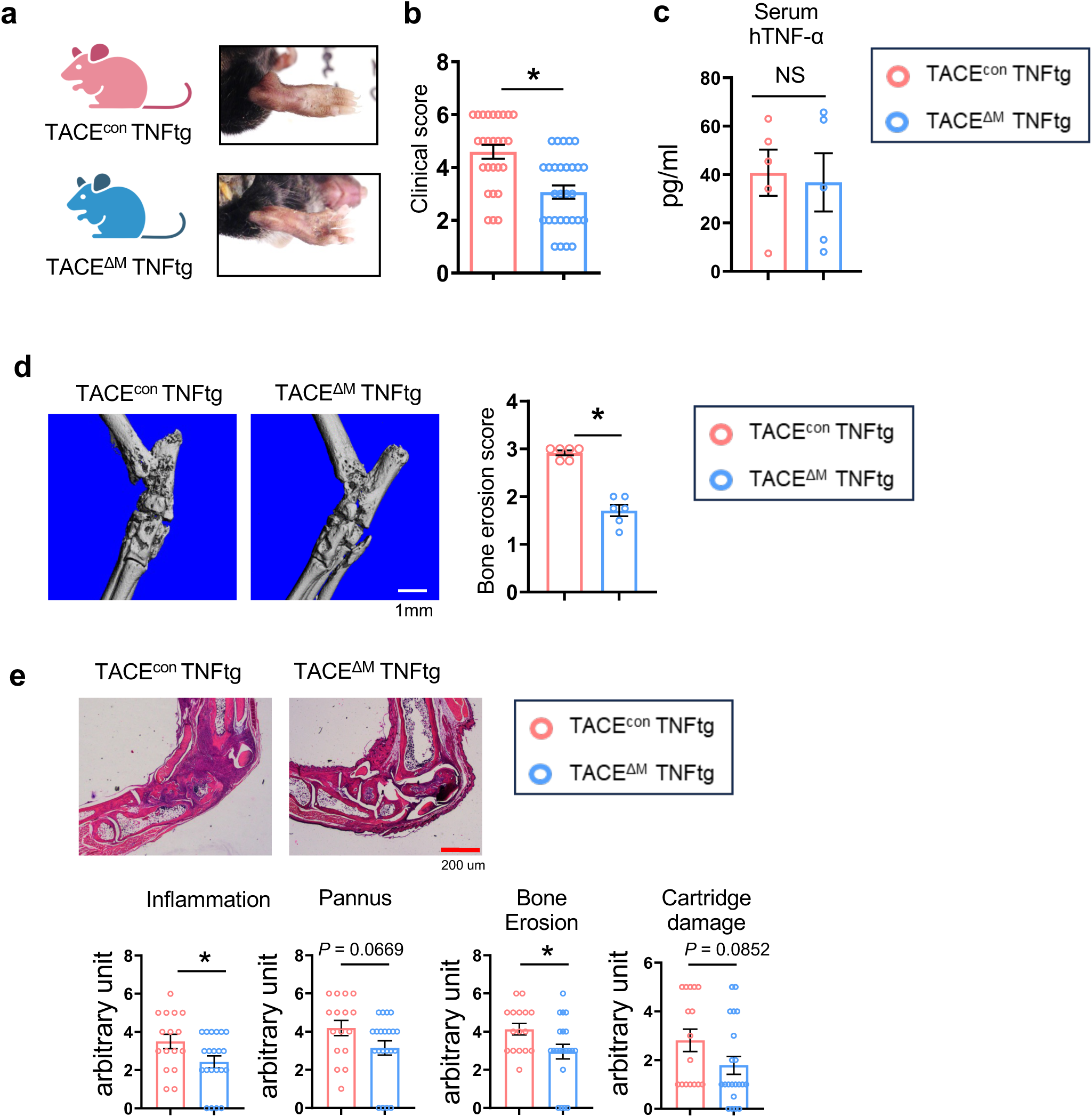
TACE deficiency attenuates inflammatory bone destruction. (**a**-**e**) Myeloid-specific TACE-deficient mice were crossed with TNFtg mice to generate TNFtg mice lacking TACE in myeloid cells (TACE^ΔM^ TNFtg). (**a**) Representative images of ankles from TACE^con^ TNFtg and TACE^ΔM^ TNFtg mice at 12 weeks of age. (**b**) Clinical arthritis scores of TACE^con^ TNFtg(n=27) and TACE^ΔM^ TNFtg(n=29). (**c**) Serum hTNF-α levels were measured by ELISA in TACE TNFtg and TACE TNFtg mice(n =5). (**d**) Bone erosion was assessed by micro-CT. Left: representative images; Right: quantification of bone erosion scores(n =6). (**e**) H&E staining of tarsal bones from hind paws of TACE TNFtg(n=16) and TACE TNFtg(n=21) mice. Upper: representative images; Lower: histological analysis of tarsal bones. White scale bar, 1 mm; red scale bar, 200 μm. All data are presented as mean ± SEM. n.s., not significant. *p < 0.05; **p < 0.01 by two-tailed, unpaired t-test (**b**, **c**, **d**, **e**).

Patients with RA are at a higher risk of osteoporosis (systemic bone loss) than other individuals without RA^39^. Accordingly, to evaluate systemic bone loss, we examined the bone phenotype through micro-CT analysis of the femurs to determine the effect of TACE deficiency on systemic bone loss using 12-week-old TACE^con^ TNFtg and TACE^ΔM^ TNFtg male mice. The absence of TACE in myeloid cells significantly attenuated the systemic bone loss compared to TACE^con^ TNFtg, which was characterized by increased bone mass (BV/TV), trabecular number (Tb.N), and trabecular separation (Tb.Sp) of the femurs (Fig. 3a,b). Histomorphometry analysis of the femur showed that osteoclast numbers, osteoclast surface, and erosion surface were significantly decreased in TACE^ΔM^ TNFtg femurs compared to TACE^con^ TNFtg mice (Fig. 3c, d). Together, our findings indicate that TACE plays an important role in arthritic bone erosion and systemic bone loss.

**Fig. 3.**
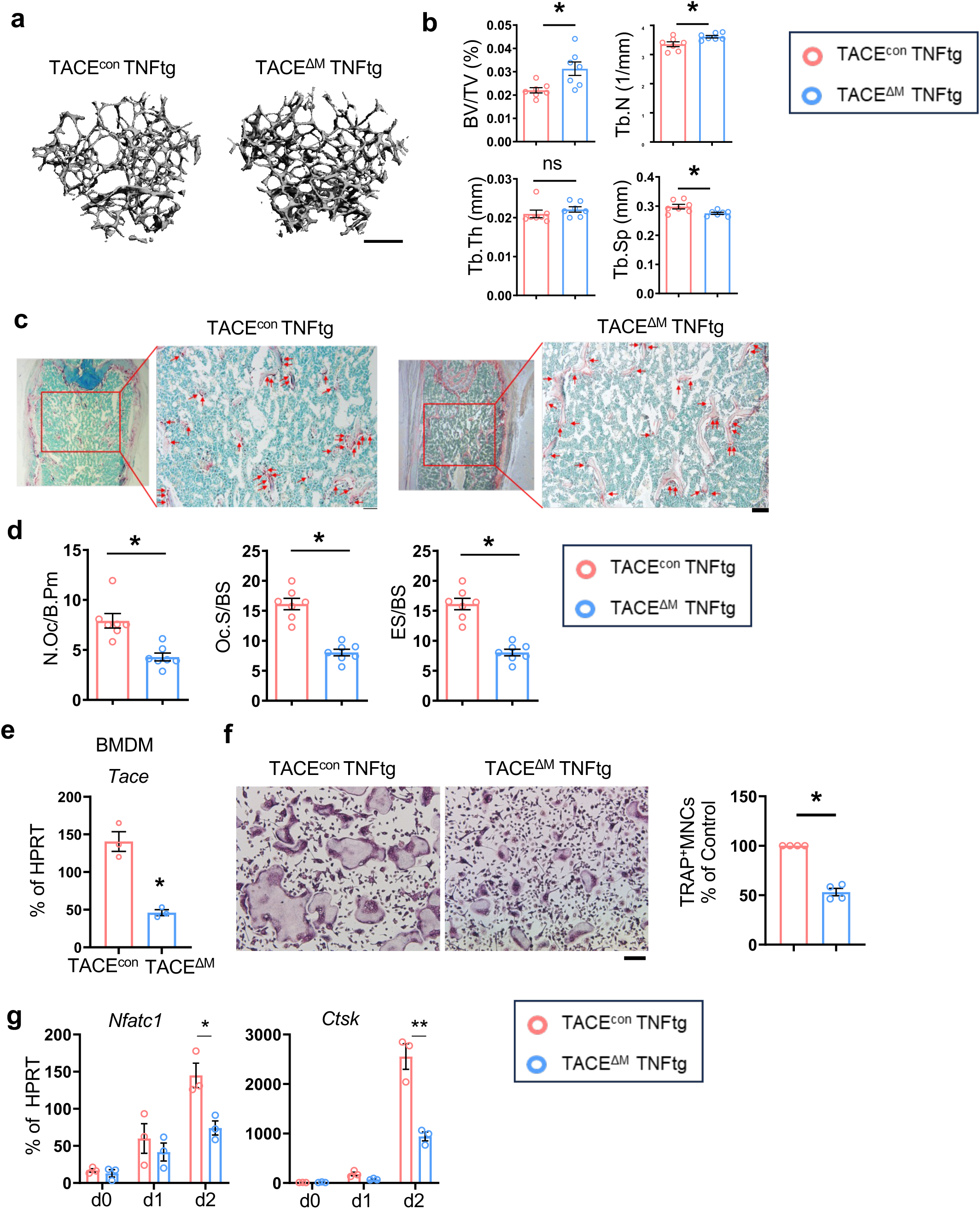
TACE positively regulates osteoclastogenesis. (**a** and **b**) Micro-CT analysis of femurs from 12-week-old TACE TNFtg and TACE TNFtg mice(n = 7). Scale bar: 0.5mm.(**a**) Representative images of the distal trabecular bone in femurs. (**b**) Analysis of bone parameters in the distal trabecular region. Bone volume/tissue volume ratio (BV/TV), trabecular thickness (Tb.Th), trabecular number (Tb.N), and trabecular separation (Tb.Sp) were quantified by micro-CT.(**c** and **d**) Histomorphometric analysis of the distal femur(n = 7). (**c**) Representative image showing TRAP-positive multinucleated osteoclasts (red). (**d**) Quantification of osteoclast number per bone surface (N.Oc/BS), osteoclast surface per bone surface (Oc.S/BS), and eroded surface per bone surface (ES/BS).(**e**-**g**) Bone marrow-derived macrophages (BMDMs) from TACE TNFtg and TACE TNFtg mice were cultured with M-CSF and RANKL for 3 days. (**e**) Efficiency of TACE knockout (KO). TACE mRNA expression was measured by RT-qPCR and normalized to HPRT. (**f**) Osteoclastogenesis assay. Left: representative images of TRAP-stained cells. Right: quantification of TRAP-positive multinucleated cells as a percentage relative to TACE TNFtg cells. Scale bar: 100 µm (**g**) mRNA expression levels of NFATc1 and CTSK were measured by RT-qPCR. All data are presented as mean ± SEM. n.s., not significant. *p < 0.05; **p < 0.01 by two-tailed, unpaired *t*-test (**b**, **d**, **e**, **f**) or one-way ANOVA with *post hoc* Tukey’s test (**g**). Data represent at least three independent experiments.

Since TACE is deleted only in myeloid cells of TACE^ΔM^ TNFtg mice, we sought to test if TACE deficiency could directly control osteoclasts. We treated BMDMs from TACE^con^ TNFtg and TACE^ΔM^ TNFtg mice with M-CSF, RANKL, and/or TNF. TACE expression was significantly diminished in TACE^ΔM^ TNFtg mice compared to control TNFtg mice (Fig. 3e). Consistent with the *in vivo* bone phenotype and the data from human cells, the number of TRAP-positive multinuclear osteoclasts was decreased in cells from TACE^ΔM^ TNFtg mice relative to TACE^con^ TNFtg cells (Fig. 3f and Supplementary Fig. 3a,b). Accordingly, the expression of osteoclast marker genes, such as *Ctsk* and *Nfatc1,* was suppressed in BMDMs from TACE^ΔM^ TNFtg mice, compared to BMDMs from TACE^con^ TNFtg mice (Fig. 3g and Supplementary Fig. 3c). Since serum TNF levels were high and cells were primed by TNF in TNFtg mice, an osteoclastogenesis assay was also performed in myeloid-specific TACE-deficient cells under a non-TNFtg background. The efficiency of deletion in TACE^ΔM^ mice was ∼60% (Supplementary Fig. 3d). Osteoclastogenesis in TACE^ΔM^ cells was significantly diminished relative to cells from TACE^con^ mice (Supplementary Fig. 3e). Taken together, our results demonstrate that TACE positively regulates osteoclastogenesis.

### TACE inhibits RANKL-induced IRF3 activation

To gain insight into the mechanisms by which TACE regulates osteoclastogenesis, we first investigate the effect of TACE on RANKL signaling pathways. The proximal RANKL signaling and p65 activation increased in TACE-deficient cells compared to control cells (Fig. 4a, b). However, NFATc1 activation, an indicator of distal RANKL signaling, was significantly diminished in TACE-deficient cells (Fig. 4c). This paradoxical observation led us to investigate the downstream pathways regulated by TACE. We performed an unbiased transcriptomic analysis using RNA-seq to identify genes whose expression was affected by TACE in the early phases of the RANKL response. Cells from TACE^con^ and TACE^ΔM^ mice were treated with M-CSF and RANKL for two days (Fig. 4d). TACE deficiency affected 157 genes (differentially expressed genes (DEGs), 2-fold changes, *p*-value <0.05); 118 genes were upregulated, and 37 genes were downregulated in TACE-deficient cells (Fig. 4e). We performed gene set enrichment analysis (GSEA)^40, 41^, the pathway broadly testing for the enrichment of well-defined gene sets from the comprehensive Molecular Signature Data Base v5.1. We found that genes involved in the IFN-α responses and the Janus kinase/signal transducer and activator of transcription (JAK/STAT) signaling pathway were significantly enriched in TACE-deficient cells compared to control cells (Fig. 4f, g). Cxcl10 and Irf7, genes in Hallmark_IFN-α response, were significantly increased in TACE^ΔM^ cells compared to TACE^con^ cells (Fig. 4h). Since negative feedback interactions of type I IFNs on RANKL-induced osteoclastogenesis have been well established ^42^, we wished to test how TACE regulates RANKL-induced interferon-stimulated genes (ISGs). To determine the mechanism, we first investigated whether TACE deficiency modulates the type I IFN signaling machinery. BMDMs from TACE^con^ and TACE^ΔM^ mice were stimulated with IFN-β; STAT1/2 activation was comparable between TACE^con^ and TACE^ΔM^ cells, suggesting that the type I IFN signaling cascades are unaffected by TACE deficiency (Supplementary Fig. 4a). We then performed upstream regulator analysis using ingenuity pathway analysis and found that Cited2, Stat1, Zbtb10, Stat6 and interferon regulatory factor 3 (IRF3) were the top upstream regulators of TACE-dependent DEGs (Fig. 4i). Among these regulators, IRF3 is a well-known inducer of type I IFN production and a key player in antiviral responses^43^. However, the role of IRF3 in osteoclasts has not been directly tested; we examined whether IRF3 is activated during osteoclastogenesis. When IRF3 was activated, IRF3 translocated to the nucleus. IRF3 was observed in the nuclear extracts three hours after RANKL stimulation and peaked one day after RANKL stimulation (Fig. 4j and Supplementary Fig. 4b). Intriguingly, IRF3 activation was further induced in TACE^ΔM^ cells compared to control cells (Fig. 4k). Our results suggested that TACE modulates RANKL-induced IRF3 activation.

**Fig. 4.**
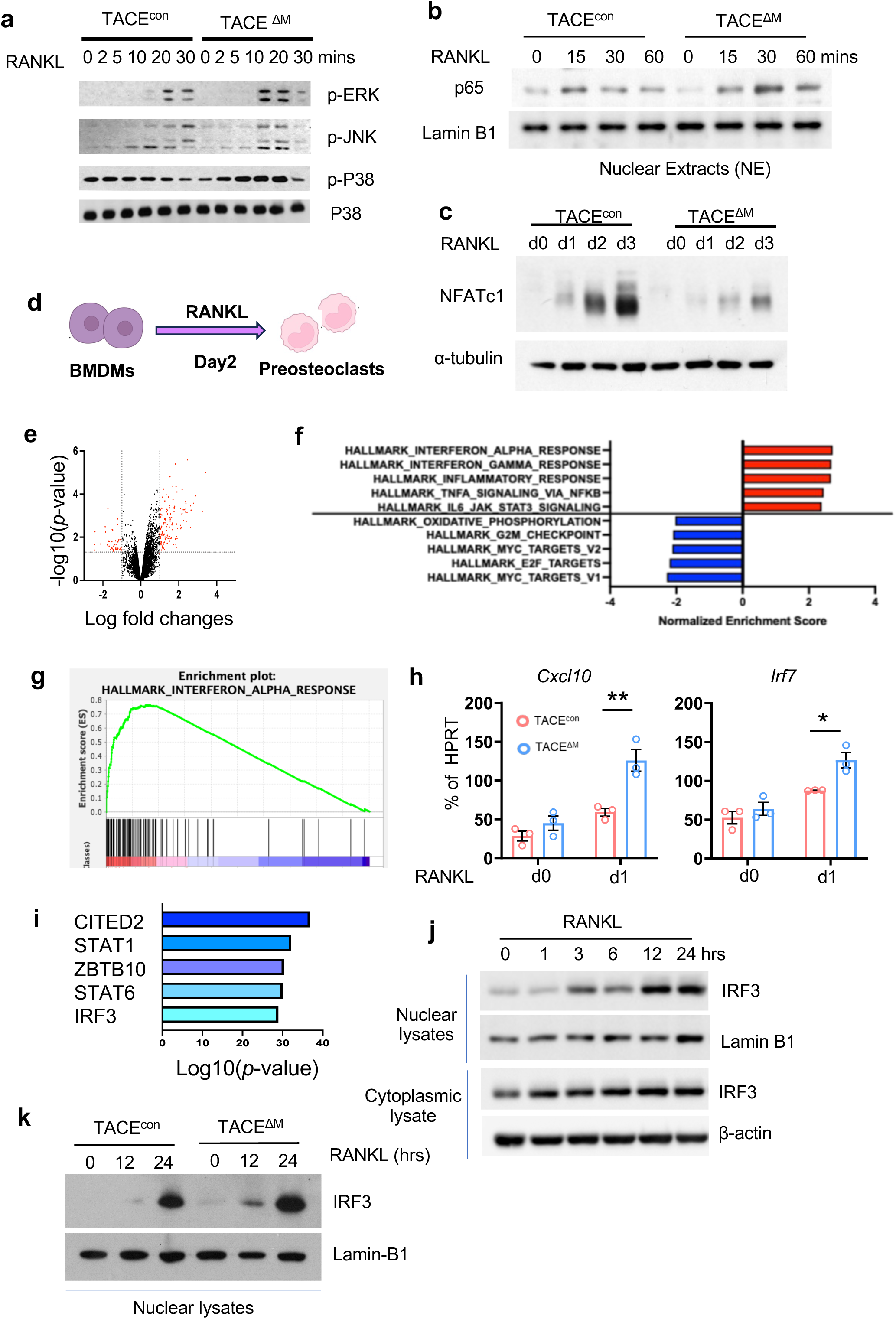
TACE promotes NFATc1 and inhibits RANKL-induced IRF3 activation. TACE ^f/f^ mice were crossed with myeloid-specific LysM-Cre mice to generate mice lacking TACE in myeloid cells (TACE^ΔM^). (**a** and **b**) BMDMs from TACE^con^ and TACE^ΔM^ mice were stimulated with RANKL (50 ng/ml) for the indicated times. (**a**) Immunoblot analysis using antibodies against phosphorylated ERK (p-ERK), JNK (p-JNK), or p38. (**b**) Nuclear p65 activity was assessed by immunoblot analysis. Lamin B1 was used as a control for nuclear fractions. (**c**) Immunoblot analysis of NFATc1 and α-tubulin during osteoclastogenesis. (**d**-**i**) BMDMs from TACE and TACE mice were treated with M-CSF and RANKL for two days. Total RNA was analyzed by bulk RNA-seq. (**d**) Schematic diagram of the experimental conditions representing the early phase of RANKL stimulation. (**e**) Volcano plot showing 39 genes upregulated and 118 genes downregulated in TACE cells (|log2 FC| ≥ 2, p < 0.05). (**f**) Gene Ontology (GO) enrichment analysis of the upregulated genes in TACE cells. (**g**) Hallmark pathway analysis of the upregulated genes in TACE cells. (**h**) Expression of IFN-dependent genes CXCL10 and IRF7 was measured by RT-qPCR. (**i**) Upstream regulator analysis was performed using Ingenuity Pathway Analysis (IPA). (**j**) BMDMs from WT mice were stimulated with RANKL (50 ng/ml) for the indicated times. Nuclear and cytosolic proteins were extracted and analyzed by immunoblotting with IRF3, Lamin B1, or α-tubulin antibodies. (**k**) BMDMs from TACE^con^ mice were stimulated with RANKL (50 ng/ml) for the indicated times. Nuclear fractions were analyzed by immunoblotting with IRF3 and Lamin B1 antibodies. All data are presented as mean ± SEM. *p < 0.05; **p < 0.01 by one-way ANOVA with *post hoc* Tukey’s test (**h**). Data represent at least three independent experiments.

### IRF3 is a negative regulator of osteoclastogenesis

To determine the effect of IRF3 on bone homeostasis, we performed micro-CT analysis of IRF3-deficient mice and wild-type mice. IRF3-deficient mice exhibited decreased trabecular bone density compared to wild-type mice (Supplementary Fig. 5a, b), while cortical bone density was comparable between IRF3-deficient mice and wild-type mice (Supplementary Fig. 5c, d). To test the direct role of IRF3 in osteoclastogenesis, bone marrow cells from control and IRF3-deficient mice were differentiated into BMDMs and then cultured with M-CSF and RANKL for three days to differentiate into osteoclasts. IRF3 expression was significantly lower in IRF3-deficient cells compared to control cells (Fig. 5a). Osteoclast formation was significantly enhanced in IRF3-deficient cells compared to control cells (Fig. 5b). To corroborate our findings, we overexpressed human IRF3 using adenoviral particles encoding IRF3 or GFP in wild-type BMDMs. IRF3 expression was significantly increased in IRF3 transduced cells (IRF3 OE) compared to GFP transduced cells (GFP) (Fig. 5c). Osteoclast formation was diminished by ectopic expression of IRF3 (Fig. 5d). To determine if TACE is an upstream regulator of RANKL-induced IRF3 activation, TACE expression was knocked down using siRNAs in WT and IRF3-deficient mice. TACE expression was sufficiently diminished by 80-90% by the TACE siRNA relative to the control siRNA in both the WT and IRF3-deficient cells (Fig. 5e). Consistently, suppression of TACE in the WT cells decreased osteoclast formation by about 50% compared to the control siRNA condition. However, enhanced osteoclastogenesis in the IRF3-deficient cells was unaffected by the abrogation of TACE expression (Fig. 5f), suggesting that TACE is an upstream regulator of RANKL-induced IRF3 activation. Our findings suggest that IRF3 negatively reprograms RANKL-induced osteoclast differentiation.

**Fig. 5.**
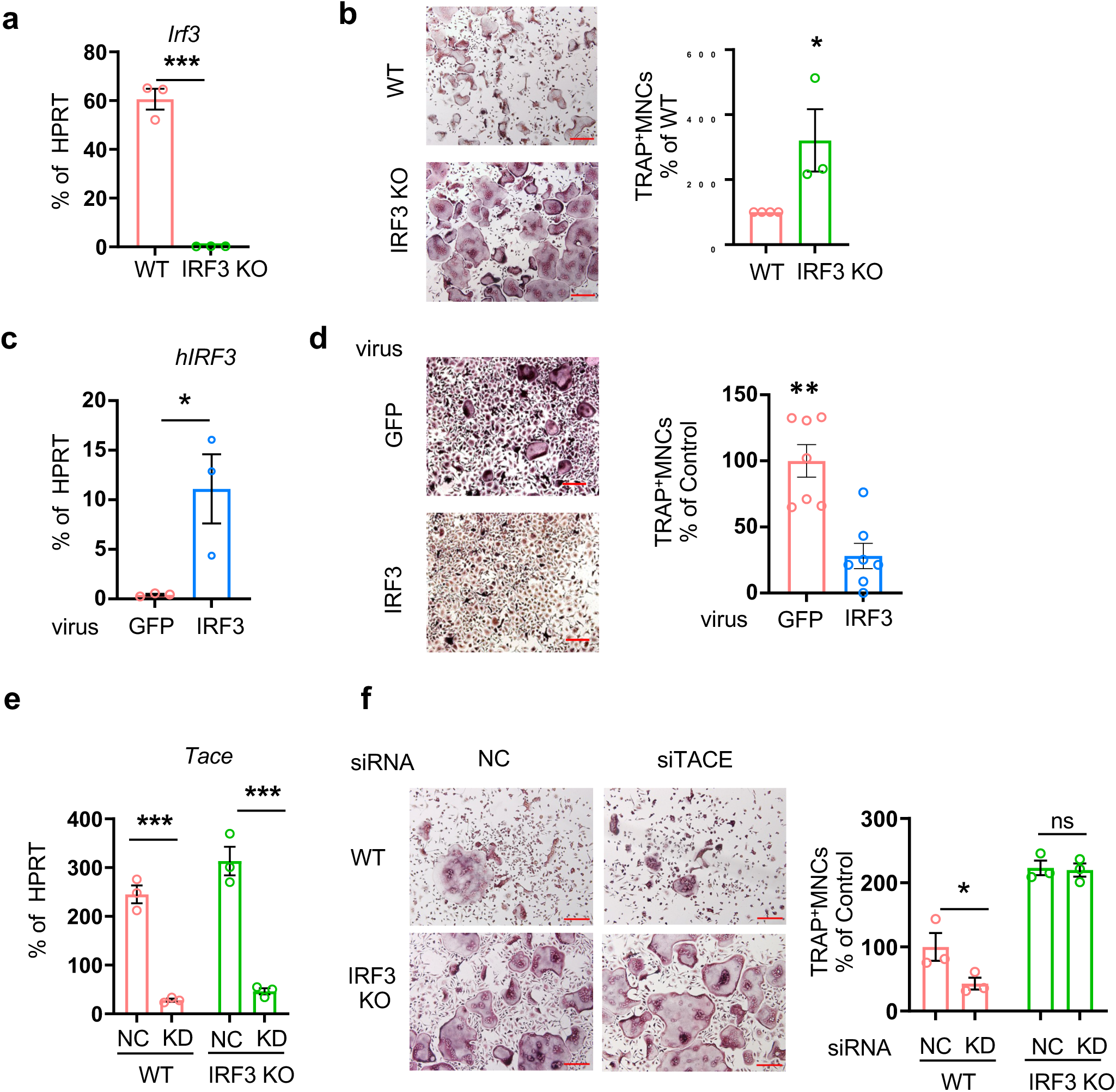
IRF3 is a negative regulator of osteoclastogenesis. (**a** and **b**) BMDMs from WT and IRF3 KO mice were cultured with M-CSF and RANKL for 3 days(n=3). (**a**) Efficiency of IRF3 knockout (KO). IRF3 mRNA expression was measured by RT-qPCR and normalized to HPRT. (**b**) Osteoclastogenesis assay. Left: representative images of TRAP-stained cells. Right: quantification of TRAP-positive multinucleated cells as a percentage relative to WT cells. (**c** and **d**) BMDM from WT mice were transduced with either control-GFP or hIRF3-expressing adenovirus and cultured with M-CSF (20 ng/ml) and RANKL (40 ng/ml) for 3 days(n=3). (**c**) Efficiency of hIRF3. hIRF3 mRNA expression was measured by RT-qPCR and normalized to HPRT (**d**) Osteoclastogenesis assay. Left: TRAP staining of representative images. Scale bar: 100 µm. Right: Quantification of TRAP-positive multinucleated cells. (**e** and **f**) BMDMs from WT and IRF3 KO mice were nucleofected with either control (CTL) or TACE-targeting siRNAs and cultured with M-CSF. (**e**) Knockdown (KD) efficiency. TACE mRNA levels were measured by qPCR and normalized to HPRT. (**f**) Osteoclastogenesis assay. Left: representative images of TRAP-stained cells. Right: quantification of TRAP-positive multinucleated cells as a percentage relative to WT-NC cells. All data are presented as mean ± SEM. n.s., not significant. Scale bar: 100 µm. *p < 0.05; **p < 0.01 by two-tailed, unpaired t-test (**b**, **c**, **d**) or one-way ANOVA with post hoc Tukey’s test (**a**, **e**, **f**). Data represent at least three independent experiments.

### IRF3 reprograms the response to RANKL in a type I IFN-dependent and independent manner

To understand how the TACE-IRF3 axis functions in osteoclastogenesis, we performed an unbiased transcriptomic analysis using RNA-seq in wild-type and IRF3-deficient cells. BMDMs from WT and IRF3-deficient mice were cultured with M-CSF and RANKL for one day. 954 differentially expressed genes (DEGs, 2 fold changes, *p*-value <0.05) between control and IRF3-deficient cells were identified; 692 genes were upregulated, and 262 genes were downregulated in IRF3-deficient cells (Fig. 6a). We performed gene set enrichment analysis (GSEA)^40, 41^. As expected, genes in Hallmark_interferon_alpha_response were downregulated in IRF3 deficient cells (Fig. 6b, c). We also measured the expression of Cxcl10 and Irf7, key ISGs, by RT-qPCR. IRF3 deficiency did not modulate immediate early induction of Cxcl10 by RANKL, while completely diminishing the expression of Cxcl10 at day one after RANKL stimulation. In contrast, IRF3 deficiency suppressed the baseline and RANKL-induced IRF7 expression (Fig. 6d). As expected, the expression of ISGs was diminished in IFNAR1-deficient cells compared to wild-type cells (Supplementary Fig. 6a), suggesting that RANKL-induced ISGs mainly depend on the type I IFN signaling pathway. MYC is a key transcription factor of osteoclastogenesis and NFATc1 expression^44^. IRF3 deficiency enhanced genes enriched in Hallmark_MYC targets_v1 and v2 (Fig. 6b). Intriguingly, NFATc1 was significantly increased in IRF3-deficient cells compared to control cells (Fig. 6e). In contrast, both protein and mRNA of NFATc1 were minimally affected by the lack of type I and II IFN signaling in IFNAR1-deficient cells and IFNα/β/γR deficient cells (Supplementary Fig. 6b-e), suggesting that IRF3 regulated RANKL-induced NFATc1 via a non-canonical pathway. To further determine the effect of type I IFNs on Nfatc1 expression, BMDMs were treated with RANKL and anti-IFNβ antibodies, an IFNβ signaling neutralizing antibody. Consistently, NFATc1 expression was minimally affected by IFNβ signaling (Supplementary Fig. 6f). Our data suggest that a non-canonical pathway of IRF3 governs the response to RANKL independently of IFN.

**Fig. 6.**
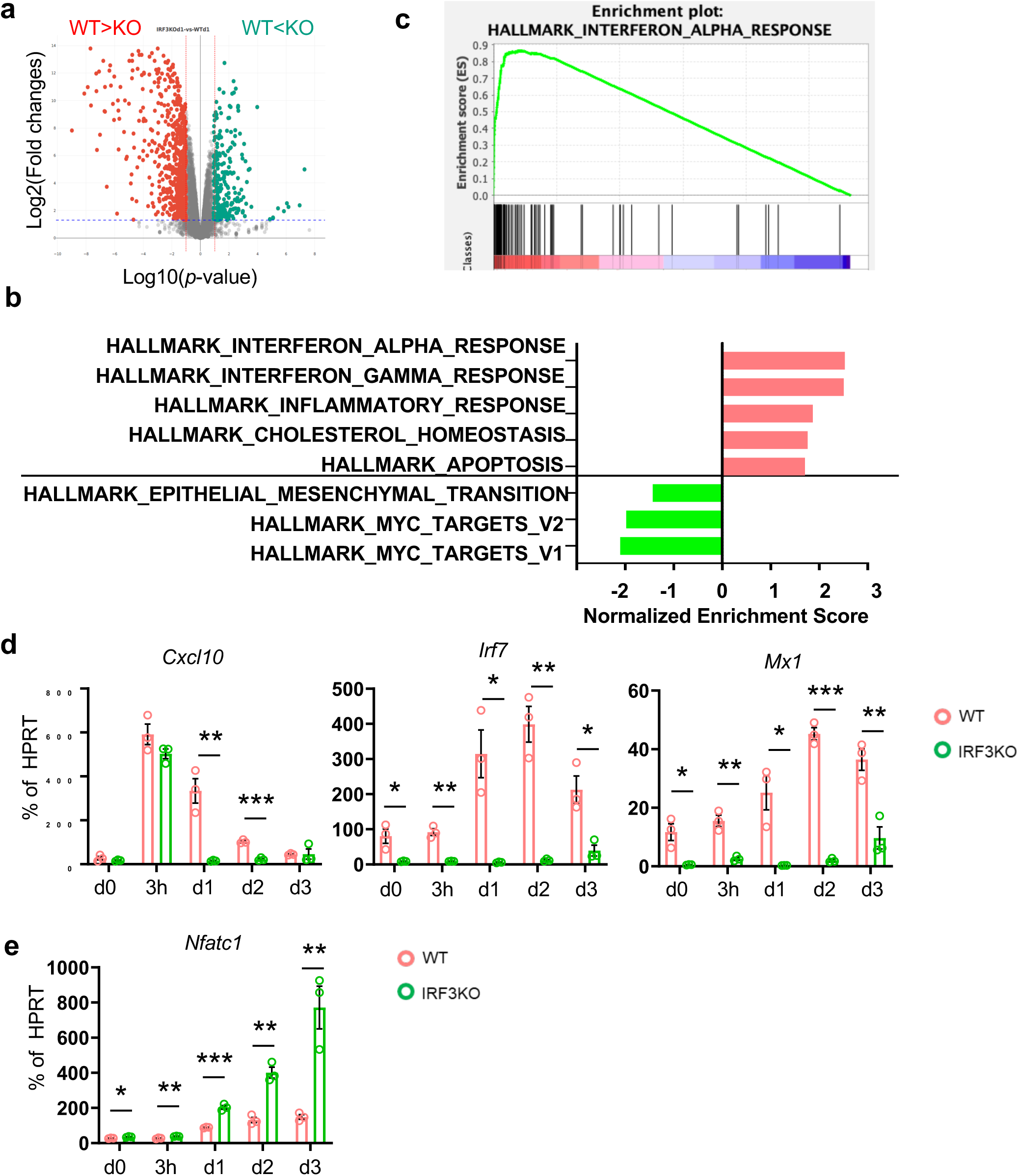
IRF3 regulates gene expression in both IFN-dependent and independent ways. (**a**-**c**) BMDMs from WT and IRF3 KO mice were treated with M-CSF and RANKL for one day. Total RNA was analyzed by bulk RNA-seq with two biological replicates. (**a**) Volcano plot showing 692 genes upregulated and 262 genes downregulated in IRF3-deficient cells (|log2 FC| ≥ 2, p < 0.05). (**b**) Gene Ontology (GO) enrichment analysis of the upregulated genes in IRF3 KO cells. (**c**) Hallmark pathway analysis of the upregulated genes in IRF3 KO cells. (**d** and **e**) BMDMs from WT and IRF3 KO mice were stimulated with RANKL (50 ng/ml) for the indicated times(n=3). (**d**) Expression of IFN-dependent genes CXCL10, IRF7, and Mx1 was measured by RT-qPCR. (**e**) Expression of NFATc1 was measured by RT-qPCR as a marker of osteoclastogenesis. All data are presented as mean ± SEM. *p < 0.05; **p < 0.01 by one-way ANOVA with post hoc Tukey’s test (**d**, **e**).

### IRF3 activation restricts the full response to RANKL in macrophages

We further investigated the regulation of IRF3-dependent genes in osteoclasts. RANKL treatment for one day induced 525 genes (Fig. 7a, Group A + B). However, increased DEGs in IRF3-deficient cells contained only part of RANKL-induced genes (96 genes, Group B). The majority of IRF3-dependent DEGs was not induced by RANKL treatment for one day in WT cells (166 genes, Group C). Among 166 genes in group C, 89 genes were upregulated on day 3 after RANKL stimulation in WT cells that led to the differentiation to mature osteoclasts, suggesting that IRF3 may delay the full activation of the expression of osteoclast marker genes. The GSEA analysis showed that in group C, genes related to TNFA_signaling_via_NF-κb were enriched (Fig. 7b). Among them, HB-EGF is a member of the EGF family of proteins and one of TACE’s substrates^45^. HB-EGF plays a crucial role in development, tissue regeneration, and cancer ^46, 47^. HB-EGF was not expressed in either wild-type or IRF3-deficient BMDMs (Fig. 7c, d), but its expression increased by RANKL stimulation and was detectable in mature WT osteoclasts (Fig. 7d). In contrast, *Hbegf* mRNA expression appeared earlier in IRF3-deficient cells on day 1 after RANKL stimulation (Fig. 7c, d), supporting the rate-limiting role of IRF3 in osteoclastogenesis. The expression of HB-EGF was comparable between control and IFNAR1-deficient cells (Fig. 7e), suggesting that IRF3 regulates RANKL-induced HB-EGF via an IFN-independent manner. HB-EGF expression was elevated in SF macrophages from RA patients (Supplementary Fig. 7a). HB-EGF+ macrophages are present in RA synovial tissues and contribute to fibroblast invasiveness and disease progression^48, 49, 50^. Single-cell RNA sequencing from the public database showed that HB-EGF was enriched within most subpopulations of RA synovium macrophages (Supplementary Fig. 7b). To determine the function of HB-EGF in osteoclastogenesis, we treated cells with RANKL and HB-EGF. HB-EGF stimulation enhanced osteoclastogenesis (Fig. 7f) and NFATc1 expression (Fig. 7g). Conversely, inhibiting EGFR signaling using erlotinib, a small molecule inhibitor for EGFR, suppressed osteoclastogenesis (Supplementary Fig. 7c). Of note, the expression of other EGFR ligands or receptors for HB-EGF, such as ErbB and ErbD, remained very low during osteoclast differentiation (data not shown). Intriguingly, HB-EGF stimulation diminished RANKL-induced IRF3 activation (Fig. 7h), suggesting that EGFR signaling may serve as a negative feedback mechanism of IRF3 activation. Our findings suggest that IRF3 activation limits the full expression of HB-EGF by RANKL during the early stage of osteoclastogenesis, while the increased expression and release of HB-EGF by TACE inhibits IRF3 activation in the later stage of osteoclast differentiation (Fig. 7i). Taken together, our results highlight the TACE-dependent cross-inhibition between IRF3 and HB-EGF as a new negative feedback loop that fine-tunes osteoclast differentiation and arthritic bone erosion.

**Fig. 7.**
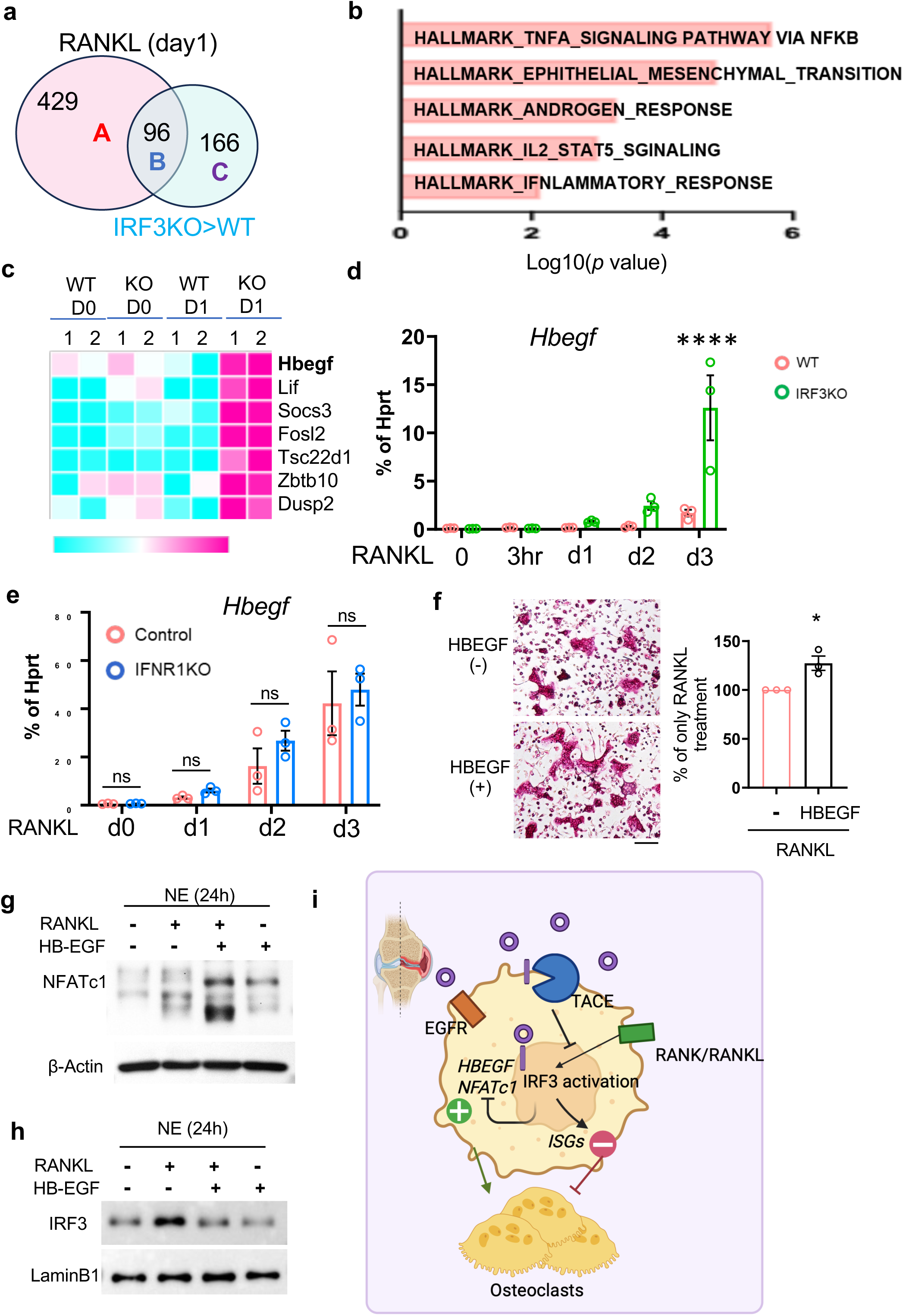
IRF3 activation suppresses HB-EGF expression in an IFN-independent manner. (**a**–**c**) BMDMs from WT and IRF3 KO mice were treated with M-CSF and RANKL for one day, and total RNA was analyzed by RNA sequencing. (**a**) Differential gene expression (DEG) analysis was performed to compare IRF3-dependent genes with those induced by RANKL. (**b**) GSEA analysis of the upregulated genes by IRF3 deficiency in Group C. (**c**) Heatmap of genes in the Hallmark_TNFA signaling pathway. (d) Expression of the Hbegf gene upon RANKL stimulation in WT and IRF3 KO BMDMs was measured by RT-qPCR. (**e**) Expression of the Hbegf gene upon RANKL stimulation in WT and IFNAR KO BMDMs was measured by RT-qPCR. (**f**–**h**) BMDMs from WT mice were cultured with M-CSF and RANKL in the presence or absence of recombinant HB-EGF (100ng/ml). (**g**) BMDMs from WT mice were stimulated with RANKL (50 ng/ml) for 24 hours. Immunoblotting was performed using NFATc1 and β-actin antibodies. (**h**) Nuclear fractions were analyzed by immunoblotting with IRF3 and Lamin B1 antibodies. (**i**) Schematic representation of TACE-dependent cross-regulation between IRF3 and HB-EGF, which fine-tunes osteoclast differentiation and arthritic bone erosion. All data are presented as mean ± SEM. Scale bar: 100 µm. *p < 0.05; ****p < 0.001 by one-way ANOVA with post hoc Tukey’s test (**d**, **e**) or two-tailed, unpaired t-test (**f**). Data represent at least three independent experiments.

## Discussion

Alleviating bone damage and suppressing synovial inflammation have emerged as potential therapeutic strategies for rheumatoid arthritis (RA)^51^. Dysregulation of osteoclast differentiation is one of the major contributors to arthritic bone erosion. However, the underlying mechanism by which hyperactive osteoclasts form in RA has not yet been fully elucidated. TACE is a metalloprotease, and its main role is to shed membrane proteins, including TNF. Therefore, many TACE functions are explained by its ability to shed TNF. Our study revealed that elevated TACE in macrophages plays a crucial role in arthritic bone erosion and osteoclastogenesis. Myeloid cell-specific TACE deficiency resulted in reduced joint inflammation and bone erosion in an inflammatory arthritis model. TACE positively regulates RANKL-induced osteoclast formation by limiting IRF3 activation. RANKL-induced IRF3 activation restricts excessive formation of osteoclasts by dual function: promotion of the expression of interferon-stimulated genes (ISGs) and suppression of osteoclast marker genes. HBEGF stimulates EGFR signaling pathways and suppresses IRF3 activation, thereby promoting RANKL-induced osteoclastogenesis. Our results reveal a new role for TACE as a rheostat in reprogramming the RANKL response by balancing the positive and negative pathways in macrophages.

Chronic inflammation reprograms macrophage cell fate and differentiation. This study shows that TACE plays a role in promoting osteoclast formation, supporting the positive effect of TACE on osteoclastogenesis in pathological and physiological conditions. Ectopic expression of TACE promotes human osteoclast differentiation, which may connect bone erosion with increased TACE in SF macrophages from RA patients. Conversely, TACE deficiency suppresses osteoclast differentiation, as observed in three different TACE-deficient lines and human macrophages. A recent study indicated that TACE deficiency does not affect the number of osteoclasts but increases their area, as shown in Vav-Cre heterozygous mice and wild-type mice^35^. The discrepancy may arise from the use of different Cre lines and the efficiency of TACE deletion. Our study shows that TACE fine-tunes RANKL-mediated macrophage differentiation toward osteoclasts. Our study uses TNF transgenic mice to compensate for the reduced release of TNF-α in TACE-deficient mice by overexpressing human TNF. Indeed, we observed comparable levels of human TNF in the serum between TACE^con^ TNFtg and TACE^ΔM^ TNFtg male mice. However, inflammation was still lower in myeloid cell-specific TACE-deficient mice compared to control mice. It is possible that systemic expression of TNF is not the only factor causing arthritic bone erosion and inflammation. Another possibility is that other TACE substrates in myeloid cells also contribute to the inflammatory responses. The shedding of substrates by TACE can significantly influence inflammatory responses and bone erosion. Our study highlights the role of TACE in regulating signaling pathways.

Our study reveals that the non-canonical IRF3 pathway controls RANKL-induced osteoclast formation. Although IRF3 is a well-known regulator of IFN-β production, its role in osteoclasts remained unclear prior to our study. We demonstrated that IRF3 targets not only the IFN-dependent pathway but also the IFN-independent pathway during osteoclast formation. IRF3 activation was elevated in TACE-deficient cells compared to control cells, suggesting that TACE suppresses RANKL-induced IRF3 activation. Despite extensive studies on RANKL-induced osteoclastogenesis, the role of RANKL in inflammatory bone loss is incompletely understood. We show the TACE’s function as a suppressor of RANKL-induced IRF3 activation during the early phase of osteoclastogenesis to promote osteoclast differentiation. However, it remains unclear how TACE inhibits IRF3 activation. IRF3 is a well-known downstream target of pattern recognition receptors, such as toll-like receptors 3 and 4, and cytosolic receptors including STING and RLRs^52^ that TACE may regulate. In immune cells, activated STING binds to TBK1, a critical downstream regulator of innate immune signaling, linking it to IRF3 and inducing the expression of type 1 interferons and ISGs^27, 28, 29^. A recent study shows that the STING pathway regulates bone remodeling^53^. However, STING deficiency minimally affects osteoclastogenesis in our system (data not shown). Further research will be necessary to fully elucidate how TACE regulates RANKL-induced IRF3 activation.

Our transcriptomic analysis suggests that IRF3 functions as a limiting step during osteoclastogenesis to prevent excessive activation. Conversely, HB-EGF inhibits IRF3 activation by activating EGFR signaling. EGFR signaling facilitates the recruitment of osteoclasts and osteoblasts to the ossification center during long bone development^54, 55^. HB-EGF is abundantly expressed in mature osteoclasts, but its expression is relatively low in macrophages. HB-EGF expression was higher in RA SF macrophages compared to control macrophages. HB-EGF+ macrophages in the synovial tissues of rheumatoid arthritis patients promote fibroblast invasiveness^49^. As HBEGF is a well-known target of TACE, our data suggests that TACE not only sheds surface HB-EGF but also regulates HB-EGF expression by modifying IRF3 activation. Thus, the TACE-IRF3-HB-EGF axis may provide another layer of regulatory mechanism for osteoclast differentiation under pathological conditions. A better understanding of the impact of the crosstalk between TACE and HB-EGF+ macrophages on hyperactive osteoclasts in RA remains an active area of research, with the precise mechanisms and interactions involved not yet fully understood. This inhibitory crosstalk between IRF3 and HB-EGF, mediated by TACE, may promote the formation of hyperactive osteoclasts in RA patients, highlighting a unique role of TACE in osteoclastogenesis and arthritic bone erosion. Our results suggest that targeting both IFN-dependent and IFN-independent pathways is crucial for regulating osteoclast formation. Therefore, understanding its regulatory mechanisms could open new avenues for therapeutic intervention in arthritic bone erosion.

## Materials and Methods

### Human CD14+ Cell Culture

Peripheral blood mononuclear cells (PBMCs) from blood leukocyte preparations purchased from the New York Blood Center were isolated by density gradient centrifugation with Ficoll (Invitrogen, Carlsbad, CA). CD14^+^ cells were obtained by isolation using anti-CD14 magnetic beads, as recommended by the manufacturer (Miltenyi Biotec, CA). Human CD14^+^ cells were cultured in α-MEM medium (Invitrogen) supplemented with 10 % fetal bovine serum (FBS, Hyclone; SH30070.03) and 1% L-glutamine with 20 ng/ml of M-CSF for 12 hours to generate osteoclast precursor cells (OCPs) The purity of monocytes was >97%, as verified by flow cytometric analysis^56^. This study was approved by the Hospital for Special Surgery’s Institutional Review Board (IRB 2019-0681).

### RNA Interference

RNA interference was performed using 0.2 nmol of three short interfering RNAs (siRNAs) targeting human TACE (Invitrogen; HSS186181) or a control siRNA (D-001810-10). These siRNAs were transfected into primary human CD14⁺ monocytes using the Amaxa Nucleofector device, set to program Y-001, with the Human Monocyte Nucleofector Kit (Amaxa), as previously described^57^.

### Adenovirus Transduction

For adenoviral transduction, recombinant adenoviral particles encoding human TACE (ADV-200349) or IRF3 (ADV-212405) and control (1060) adenoviral particles encoding green fluorescent protein (Ad-CMV-GFP) were purchased from Vector Biolabs (Malvern, PA, USA). Human CD14⁺ cells were cultured at a density of 1.5 × 10⁶ cells per mL for 6 days in α-MEM medium supplemented with 10% FBS, 1% L-glutamine, and M-CSF (40 ng/mL). After incubation, the cells were washed and transferred to low-serum medium (2% FBS) containing 20 ng/mL of M-CSF and adenoviral particles (100 particles per cell) overnight. Following transduction, the cells were used for experiments, and some were subsequently replaced for osteoclastogenesis studies^58^.

### Osteoclast formation

For human osteoclastogenesis assays, cells were added to 96 well plates in triplicate at a seeding density of 5×10^4^ cells per well. Osteoclast precursors were incubated with 20 ng/ml of M-CSF and 40 ng/ml of human soluble RANKL up to 5 days in α-MEM supplemented with 10 % FBS and 1% L-glutamine. Cytokines were replenished every 3 days. On each day, cells were fixed and stained for TRAP using the Acid Phosphatase Leukocyte diagnostic kit (Sigma; 387A) as recommended by the manufacturer. Multinucleated (greater than 3 nuclei), TRAP-positive osteoclasts were counted in triplicate wells. For mouse osteoclastogenesis, bone marrow (BM) cells were flushed from femurs of mice and cultured with murine M-CSF (20 ng/ml) on petri dishes in α-MEM supplemented with 10% FBS, 1% antibiotics and 1% L-glutamine after lysis of RBCs using ACK lysis buffer (Gibco). Then, the non-adherent cell population was recovered the next day and cultured with M-CSF-containing conditional medium (CM) for three additional days. We defined this cell population as mouse OCPs. For murine osteoclastogenesis assays, we plated 2×10^4^ OCPs per well in triplicate wells on a 96 well plate and added M-CSF (20 ng/ml) and RANKL (50 ng/ml) up to 4 days, with exchange of fresh media every 3 days. All cell-cultures were performed by a modification of previously published method^56^.

### RNA preparation and real-time PCR

DNA-free RNA was obtained using the RNeasy Mini Kit from QIAGEN with DNase treatment, and 0.5 μg of total RNA was reverse transcribed using a First Strand cDNA Synthesis kit (Fermentas, Hanover, MD). Real time PCR was performed in triplicate using the iCycler iQ thermal cycler and detection system (Applied Biosystems, Carlsbad, CA) following the manufacturer’s protocols. Primer sequences are provided in the Table S1.

### Immunoblot

Whole cell extracts were prepared by lysis in buffer containing 1x Lamin sample buffer (Bio-rad) and 2-Mercaptoethanol (Sigma). The cell membrane-permeable protease inhibitor, Pefablock (1 mM), was added immediately prior to harvest cells. The membrane proteins were extracted with Mem-PER™ Plus Membrane Protein Extraction Kit (Thermofisher scientific; 89842) according to manufacturer’s instructions. To extracts of nucleus protein, cells were incubated in buffer A (10 mM Hepes, pH 7.9, 1.5 mM MgCl2, 10 mM KCl, 0.1mM EDTA, 0.1mM EGTA, proteinase inhibitor cocktail (Complete, Roche) and 1mM DTT) for 15 min at 4°C. NP-40 was added to a final concentration of 0.5%. Nuclear samples were collected by centrifugation (5000 rpm, 5 min). The pellets were lysed by Bioruptor-ultrasonicator (UCD400, Diagenode) in buffer B (20 mM Hepes, pH 7.9, 0.4M NaCl, 10 mM KCl, 1mM EDTA, 1mM EGTA, 10% Glycerol, proteinase inhibitor cocktail and 1mM DTT) and collected the supernatant by centrifugation (12,000 rpm, 10 min). The protein concentration of nuclear extracts was quantitated using the Bradford assay (Bio-Rad; 5000001). For immunoblot, proteins were separated on 7.5 or 10% SDS-PAGE gels, transferred to polyvinylidene difluoride membranes (PVDF, Millipore; ISEQ00010), and detected by antibodies as listed in the Fig. legends.

### In vivo animal studies

All animals were maintained in a pathogen-free environment and were handled according to protocols approved by the Institutional Animal Care and Use committee at Weill Cornell Medical College(2015-0062). All animals were randomly assigned into experimental groups. 8 to 12 weeks-old mice were used. TNF transgenic mice were kindly provided by Dr. George Kollias (Biomedical Sciences Research Center "Alexander Fleming)^59^. TACE ^flox/flox^ mice and TACE null mice^60^ were kindly provided by Dr. Blobel (Hospital for Special Surgery, New York, NY). IRF3 knockout mice were obtained through collaboration with the Liang Deng lab (Memorial Sloan Kettering). TACE ^flox/flox^ mice was crossed to LysM-Cre transgenic mice, which were purchased from The Jackson Laboratory (Bar Harbor, ME). The TACE ^flox/flox^/ LysM-Cre (TACE^ΔM^) generated as lacking TACE in myeloid/osteoclast lineage. TACE^ΔM^ were crossed with human TNF transgenic mouse (T197)^59^. This mouse was kindly provided by Dr. Kalliolias. IFNR DKO (IFNAR1/IFNGR1 double-deficient) mice were kindly provided by Dr. Yong-Suk JANG (Jeonbuk National University, South Korea)^61^. Finally, we generated TACE^ΔM^ (hTNF)tg (mice. Littermate TACE ^+/+ and^ ^+/-^ (TACE^con^ tg) mice were used as controls. All experiments used 12 weeks-old mice. The development of arthritis was measured by clinical scoring of the ankle joints in blinded manner. For each animal, the scoring was calculated as the sum of the measurements of both ankles of hind paws (0 = Normal paw, 1 = Slight inflamed and swollen paw, 2 = More than inflamed and swollen, or Mild swelling of ankle, 3 = Very inflamed and swollen paw or ankylosed paw. The erosion levels of the ankles of hind paws were assessed using micro-computed tomography (micro-CT) and manually scored by three independent observers in a blinded manner. Erosion was graded on a 4-point scale (Score 0: no detectable erosion; Score 1: small, isolated pits on the cortical surface; Score 2: larger defects visibly altering the bone surface contour; Score 3: extensive cortical bone destruction affecting joint integrity). The final erosion score for each sample was determined by calculating the mean of the three independent measurements^62^

### Flow Cytometry

Isolated splenocytes from TACE^con^, TACE^ΔM^ tg, and TACE^con^tg mice were treated with ACK lysis buffer (Sigma-Aldrich) to remove erythrocytes. To block nonspecific staining, cells were incubated with anti-mouse CD16/CD32 antibodies for 10 minutes on ice. Cells were then stained with B220 antibody (for B cells, BioLegend) and CD3 antibody (for T cells, BioLegend) at 4°C for 30 minutes in the dark. After washing, the stained cells were analyzed using flow cytometry. A gating strategy was applied to distinguish B cells (B220⁺) and T cells (CD3⁺) based on fluorescence intensity. Flow cytometry data were analyzed using FlowJo software.

### Enzyme-linked immunosorbent assay (ELISA)

Human TNF⍺ or mouse TNF⍺ in serum from TACE^ΔM^ TNF-Ttg and TACE^con^ TNF-tg mice were measured using ELISA kit (R&D Systems; DY210 or DY410) according to the manufacturer’s instructions

### Micro-Computed Tomography (μ-CT)

μ-CT analysis was performed as described previously^63^, and all femur samples were included in the analysis conducted in a blinded manner. For μCT analysis, a Scanco Medical μCT 35 system with an isotropic voxel size of 7 μm was used to image the distal femur. Scans were conducted in 70% ethanol and used an X-ray tube potential of 55 kVp, an X-ray intensity of 0.145 mA, and an integration time of 600 ms. For analysis of femoral bone mass, a region of trabecular bone 2.1 mm wide was contoured, starting 280 μm from the proximal end of the distal femoral growth plate. A total of 250 slices were read in each sample. Femoral trabecular bone was thresholded at 211 per mille. Femoral cortical bone was thresholded at 350 per mille.

### Histomorphometry analysis

Hematoxylin and Eosin (H&E)-stained sections of each animal’s ankle were analyzed, and the total arthritis severity was determined by summing the measurements from both wrists and both ankles, with scoring conducted blindly by three independent investigators to ensure unbiased evaluation. Using a previously established modified semiquantitative scoring system^64^, arthritis severity was assessed based on pannus formation, inflammatory cell infiltration, bone erosion, and cartilage degradation. The histology of H&E staining was examined by an HSS research pathologist. Histomorphometry samples were obtained from the femur bone of TACE^con^ tg and TACE^ΔM^ tg mice. Bone histomorphometric analysis was performed in a blinded, nonbiased manner using a computerized semi-automated system (Osteomeasure, TN) with light microscopy. Femurs were fixed in 4% paraformaldehyde for 2 days, decalcified with 14% neutral buffered EDTA (Sigma-Aldrich), and embedded in paraffin. The quantification of osteoclasts was performed in paraffin embedded tissues that were stained for TRAP and counterstained with methyl green^13^. Osteoclasts were identified as multinucleated TRAP-positive cells adjacent to bone surfaces. The measurement terminology and units used for histomorphometric analysis were those recommended by the Nomenclature Committee of the American Society for Bone and Mineral Research^65^.

### RNA Sequencing

The pre-processed data from RNAs of RA synovial macrophages were obtained from GSE97779^66^. For bulk RNA-seq experiments for TACE-deficient or IRF3-deficient cells, total RNA was purified with the RNeasy Mini kit (Qiagen) and treated on-column with DNase to eliminate genomic DNA. Poly-adenylated transcripts were captured and indexed libraries were generated using Illumina TruSeq reagents. Library integrity was confirmed on an Agilent 2100 Bioanalyzer, and all libraries satisfied quality requirements. Paired-end sequencing (50 bp × 2, ∼75 million read pairs per sample) was carried out on an Illumina HiSeq 2500 at the Weill Cornell Genomics Resources Core Facility. Reads were aligned to the mouse reference genome (mm10) with TopHat v2.0.11, and transcript abundance was estimated with Cufflinks v2.2.0, reported as FPKM (fragments per kilobase of transcript per million mapped reads). Transcripts with FPKM < 5 were excluded from downstream analyses. Differential expression was assessed with Gene Set Enrichment Analysis (GSEA; https://www.gsea-msigdb.org/gsea/), using the normalized enrichment score (NES) as the primary statistic. All bioinformatic analyses were based on two independent biological replicates.

### Single-cell RNA-sequencing

The pre-processed gene-cell matrices of the scRNA seq data^67, 68, 69^ were obtained from EMBL’s European Bioinformatics Institute (EMBL-EBI) database (Array Express: E-MTAB-8322); the data were obtained from and reanalyzed following the methods of a previous study^70^. The data were analyzed using the Seurat (v4.1.0) R package for quality control, filtering, and downstream analysis as previously described. Cells with a mitochondrial gene percentage greater than 1%, or fewer than 200 genes or more than 6,000 genes, were filtered out. Some samples were filtered more strictly, as indicated in the reference paper. Log normalization methods were used for normalizing the gene expression data. Clustering was performed using the shared nearest neighbor (SNN) algorithm. Cell types were identified as indicated in the reference article^70^. The expression of the HBEGF gene was determined using the Wilcoxon rank-sum test.

### Statistical analysis

In all experiments, data are presented as mean ± SEM if not stated otherwise. All statistical analyses were performed with GraphPad Prism 8.0 software using the two-tailed, paired or unpaired t-test (two conditions), One-way, or Two-way ANOVA for multiple comparisons (more than two conditions) with a post hoc Tukey test. P < 0.05 (*) was taken as statistically significant. Sample sizes were chosen according to standard guidelines. Number of animals was indicated as ‘‘n.’’

## Data availability

The bulk RNA-seq datasets generated by the authors as part of this study are deposited in the Gene Expression Omnibus database (GSE310585). Source data are provided.

## Patient and public involvement

Patients or the public were not involved in the design, conduct, reporting, or dissemination plans of our research.

## Ethics approval

Weill Cornell IACUC approved all animal protocols (2015-0062). Human materials were approved by the Hospital for Special Surgery’s Institutional Review Board (IRB 2019-0681).

## Acknowledgements

We thank Drs. Lionel Ivashkiv and Carl Blobel for helpful discussion. We thank Weill Cornell Medicine Genomics Resource Core for RNA sequencing and the HSS Micro-CT core for micro-CT scanning. This work was supported by the National Institute of Arthritis and Musculoskeletal and Skin Diseases (NIAMS) of NIH under Award Number AR073156, AR08176, and AR 083374 (to K.H. P.-M.), and by the support for the Rosensweig Genomics Center from The Tow Foundation (to K.H. P.-M.).

## Contributions

All authors agree to be accountable for all aspects of the work. Reviews were done by all authors. Methodology was undertaken by all authors. Conceptualization was done by S.M. and K.-H. P.-K. Computational analysis was undertaken by B.O., D.O., G.K, and K.K. Data curation was performed by S.M., B.O., S.Y., A.U., A.S., K.K., and W.K.. TP performed histological analysis. Y.Y. and L.D. provided mice and data interpretation. Writing of the original draft was done by S.M. Editing was done by S.M., B.O., S.Y., and K.-H. P.-K. K.-H. P.-K. oversaw the study.

## References

1. Gravallese EM, Firestein GS. Rheumatoid Arthritis - Common Origins, Divergent Mechanisms. N Engl J Med 388, 529–542 (2023).

2. Goldring SR. Pathogenesis of bone and cartilage destruction in rheumatoid arthritis. Rheumatology (Oxford) 42 **Suppl 2**, ii11–16 (2003).

3. Redlich K, Smolen JS. Inflammatory bone loss: pathogenesis and therapeutic intervention. Nat Rev Drug Discov 11, 234–250 (2012).

4. Boyle WJ, Simonet WS, Lacey DL. Osteoclast differentiation and activation. Nature 423, 337–342 (2003).

5. Takayanagi H. Osteoimmunology: shared mechanisms and crosstalk between the immune and bone systems. Nature reviews Immunology 7, 292–304 (2007).

6. Park-Min KH. Mechanisms involved in normal and pathological osteoclastogenesis. Cell Mol Life Sci 75, 2519–2528 (2018).

7. McInnes IB, Schett G. The pathogenesis of rheumatoid arthritis. N Engl J Med 365, 2205–2219 (2011).

8. Tsukasaki M, Takayanagi H. Osteoimmunology: evolving concepts in bone-immune interactions in health and disease. Nat Rev Immunol 19, 626–642 (2019).

9. Novack DV, Teitelbaum SL. The osteoclast: friend or foe? Annu Rev Pathol 3, 457–484 (2008).

10. Zhang F, et al. Defining inflammatory cell states in rheumatoid arthritis joint synovial tissues by integrating single-cell transcriptomics and mass cytometry. Nat Immunol 20, 928–942 (2019).

11. Hasegawa T, et al. Identification of a novel arthritis-associated osteoclast precursor macrophage regulated by FoxM1. Nat Immunol 20, 1631–1643 (2019).

12. Xia X, et al. Single cell immunoprofile of synovial fluid in rheumatoid arthritis with TNF/JAK inhibitor treatment. Nat Commun 16, 2152 (2025).

13. Mun SH, et al. Augmenting MNK1/2 activation by c-FMS proteolysis promotes osteoclastogenesis and arthritic bone erosion. Bone Res 9, 45 (2021).

14. Park JH, Lee NK, Lee SY. Current Understanding of RANK Signaling in Osteoclast Differentiation and Maturation. Mol Cells 40, 706–713 (2017).

15. Arai F, Miyamoto T, Ohneda O, Inada T, Sudo T, Brasel K, Miyata T, Anderson DM, Suda T Commitment and differentiation of osteoclast precursor cells by the sequential expression of c-fms and receptor activator of nuclear factor kB (RANK) receptors. J Experimental Medicine 190, 1741–1754 (1999).

16. Yasuda H, Shima N, Nakagawa N, Yamaguchi K, Kinosaki M, Mochizuki S, Tomoyasu A, Yano K, Goto M, Murakami A, Tsuda E, Morinaga T, Higashio K, Udagawa N, Takahashi N, Suda T. Osteoclast differentiation factor is a ligand for osteoprotegerin/osteoclastogenesis-inhibitory factor and is identical to TRANCE/RANKL. Proceedings of National Academy of Sciences USA 95, 3597–3602 (1998).

17. Anderson D, Maraskovsky E, Billingsley WL, Dougall WC, Tometsko ME, Roux ER, Teepe MC, Dubose RF, Cosman D, Galibert L. A homologue of the TNF receptor and its ligand enhance T-cell growth and dendritic-cell function. Nature 390, 175–179 (1997).

18. Choi Y, Arron JR, Townsend MJ. Promising bone-related therapeutic targets for rheumatoid arthritis. Nat Rev Rheumatol 5, 543–548 (2009).

19. Lam J, Takeshita S, Barker JE, Kanagawa O, Ross FP, Teitelbaum SL. TNF-alpha induces osteoclastogenesis by direct stimulation of macrophages exposed to permissive levels of RANK ligand. J Clin Invest 106, 1481–1488 (2000).

20. Kim N, et al. Osteoclast differentiation independent of the TRANCE-RANK-TRAF6 axis. J Exp Med 202, 589–595 (2005).

21. Hu Q, Zhong X, Tian H, Liao P. The Efficacy of Denosumab in Patients With Rheumatoid Arthritis: A Systematic Review and Pooled Analysis of Randomized or Matched Data. Front Immunol 12, 799575 (2021).

22. Cohen SB, et al. Denosumab treatment effects on structural damage, bone mineral density, and bone turnover in rheumatoid arthritis: a twelve-month, multicenter, randomized, double-blind, placebo-controlled, phase II clinical trial. Arthritis Rheum 58, 1299–1309 (2008).

23. Xiong Q, Zhang L, Ge W, Tang P. The roles of interferons in osteoclasts and osteoclastogenesis. Joint Bone Spine 83, 276–281 (2016).

24. Takayanagi H, Kim S, Taniguchi T. Signaling crosstalk between RANKL and interferons in osteoclast differentiation. Arthritis Res 4 **Suppl 3**, S227–232 (2002).

25. Wang C, et al. Gut microbiota and metabolites as predictors of biologics response in inflammatory bowel disease: A comprehensive systematic review. Microbiol Res 282, 127660 (2024).

26. Honda K, Taniguchi T. IRFs: master regulators of signalling by Toll-like receptors and cytosolic pattern-recognition receptors. Nat Rev Immunol 6, 644–658 (2006).

27. Kwon Y, Park OJ, Kim J, Cho JH, Yun CH, Han SH. Cyclic Dinucleotides Inhibit Osteoclast Differentiation Through STING-Mediated Interferon-beta Signaling. J Bone Miner Res 34, 1366–1375 (2019).

28. Ishikawa H, Ma Z, Barber GN. STING regulates intracellular DNA-mediated, type I interferon-dependent innate immunity. Nature 461, 788–792 (2009).

29. Samson N, Ablasser A. The cGAS-STING pathway and cancer. Nat Cancer 3, 1452–1463 (2022).

30. Scheller J, Chalaris A, Garbers C, Rose-John S. ADAM17: a molecular switch to control inflammation and tissue regeneration. Trends Immunol 32, 380–387 (2011).

31. Arribas J, Esselens C. ADAM17 as a therapeutic target in multiple diseases. Current pharmaceutical design 15, 2319–2335 (2009).

32. Lisi S, D’Amore M, Sisto M. ADAM17 at the interface between inflammation and autoimmunity. Immunol Lett 162, 159–169 (2014).

33. Issuree PD, et al. iRHOM2 is a critical pathogenic mediator of inflammatory arthritis. J Clin Invest 123, 928–932 (2013).

34. Boissy P, et al. An assessment of ADAMs in bone cells: absence of TACE activity prevents osteoclast recruitment and the formation of the marrow cavity in developing long bones. FEBS Lett 553, 257–261 (2003).

35. Babendreyer A, et al. Downregulation of the metalloproteinases ADAM10 or ADAM17 promotes osteoclast differentiation. Cell Commun Signal 22, 322 (2024).

36. Sorensen MG, et al. Characterization of osteoclasts derived from CD14+ monocytes isolated from peripheral blood. J Bone Miner Metab 25, 36–45 (2007).

37. Welsing PM, van Gestel AM, Swinkels HL, Kiemeney LA, van Riel PL. The relationship between disease activity, joint destruction, and functional capacity over the course of rheumatoid arthritis. Arthritis Rheum 44, 2009–2017 (2001).

38. Kotake S, et al. Activated human T cells directly induce osteoclastogenesis from human monocytes: possible role of T cells in bone destruction in rheumatoid arthritis patients. Arthritis Rheum 44, 1003–1012 (2001).

39. Sambrook PN, et al. Osteoporosis in rheumatoid arthritis. A monozygotic co-twin control study. Arthritis Rheum 38, 806–809 (1995).

40. Subramanian A, et al. Gene set enrichment analysis: a knowledge-based approach for interpreting genome-wide expression profiles. Proc Natl Acad Sci U S A 102, 15545–15550 (2005).

41. Mootha VK, et al. PGC-1alpha-responsive genes involved in oxidative phosphorylation are coordinately downregulated in human diabetes. Nature genetics 34, 267–273 (2003).

42. Takayanagi H, et al. RANKL maintains bone homeostasis through c-Fos-dependent induction of interferon-beta. Nature 416, 744–749 (2002).

43. Andersen J, VanScoy S, Cheng TF, Gomez D, Reich NC. IRF-3-dependent and augmented target genes during viral infection. Genes Immun 9, 168–175 (2008).

44. Bae S, et al. MYC-dependent oxidative metabolism regulates osteoclastogenesis via nuclear receptor ERRalpha. J Clin Invest 127, 2555–2568 (2017).

45. Jackson LF, et al. Defective valvulogenesis in HB-EGF and TACE-null mice is associated with aberrant BMP signaling. EMBO J 22, 2704–2716 (2003).

46. Dao DT, Anez-Bustillos L, Adam RM, Puder M, Bielenberg DR. Heparin-Binding Epidermal Growth Factor-Like Growth Factor as a Critical Mediator of Tissue Repair and Regeneration. Am J Pathol 188, 2446–2456 (2018).

47. Miller A, Hafler DA, Weiner HL. Immunotherapy in autoimmune diseases. Curr Opin Immunol 3, 936–940 (1991).

48. Alivernini S, et al. Distinct synovial tissue macrophage subsets regulate inflammation and remission in rheumatoid arthritis. Nat Med 26, 1295–1306 (2020).

49. Kuo D, et al. HBEGF(+) macrophages in rheumatoid arthritis induce fibroblast invasiveness. Sci Transl Med 11, (2019).

50. Baker KF, et al. Single-cell insights into immune dysregulation in rheumatoid arthritis flare versus drug-free remission. Nat Commun 15, 1063 (2024).

51. Schett G, Gravallese E. Bone erosion in rheumatoid arthritis: mechanisms, diagnosis and treatment. Nat Rev Rheumatol 8, 656–664 (2012).

52. Wang L, et al. The multiple roles of interferon regulatory factor family in health and disease. Signal Transduct Target Ther 9, 282 (2024).

53. MacLauchlan S, et al. STING-dependent interferon signatures restrict osteoclast differentiation and bone loss in mice. Proc Natl Acad Sci U S A 120, e2210409120 (2023).

54. Zhu J, Shimizu E, Zhang X, Partridge NC, Qin L. EGFR signaling suppresses osteoblast differentiation and inhibits expression of master osteoblastic transcription factors Runx2 and Osterix. J Cell Biochem 112, 1749–1760 (2011).

55. Wang K, Yamamoto H, Chin JR, Werb Z, Vu TH. Epidermal growth factor receptor-deficient mice have delayed primary endochondral ossification because of defective osteoclast recruitment. J Biol Chem 279, 53848–53856 (2004).

56. Park-Min KH, et al. Inhibition of osteoclastogenesis and inflammatory bone resorption by targeting BET proteins and epigenetic regulation. Nature communications 5, 5418 (2014).

57. Park-Min KH, et al. FcgammaRIII-dependent inhibition of interferon-gamma responses mediates suppressive effects of intravenous immune globulin. Immunity 26, 67–78 (2007).

58. Murata K, et al. Hypoxia-Sensitive COMMD1 Integrates Signaling and Cellular Metabolism in Human Macrophages and Suppresses Osteoclastogenesis. Immunity 47, 66–79 e65 (2017).

59. Keffer J, et al. Transgenic mice expressing human tumour necrosis factor: a predictive genetic model of arthritis. The EMBO journal 10, 4025–4031 (1991).

60. Horiuchi K, et al. Cutting edge: TNF-alpha-converting enzyme (TACE/ADAM17) inactivation in mouse myeloid cells prevents lethality from endotoxin shock. J Immunol 179, 2686–2689 (2007).

61. Baldon LVR, et al. AG129 Mice as a Comprehensive Model for the Experimental Assessment of Mosquito Vector Competence for Arboviruses. Pathogens 11, (2022).

62. Bouxsein ML, Boyd SK, Christiansen BA, Guldberg RE, Jepsen KJ, Muller R. Guidelines for assessment of bone microstructure in rodents using micro-computed tomography. J Bone Miner Res 25, 1468–1486 (2010).

63. Shim JH, et al. Schnurri-3 regulates ERK downstream of WNT signaling in osteoblasts. J Clin Invest 123, 4010–4022 (2013).

64. Buttgereit F, et al. Transgenic disruption of glucocorticoid signaling in mature osteoblasts and osteocytes attenuates K/BxN mouse serum-induced arthritis in vivo. Arthritis Rheum 60, 1998–2007 (2009).

65. Parfitt A, Drezner MK, Vlorieux FH, Kanis JA, Malluche H, Meunier PJ, Ott SM, Recker RR. Bone histomorphometry:stardization of nomenclature, symbols, and units. Report of the ASBMR Histomorphometry Nomenclature Committee. J Bone Mineral Research 2, 595–610 (1987).

66. Kang K, et al. Interferon-γ Represses M2 Gene Expression in Human Macrophages by Disassembling Enhancers Bound by the Transcription Factor MAF. Immunity 47, 235–250.e234 (2017).

67. Alivernini S, et al. Distinct synovial tissue macrophage subsets regulate inflammation and remission in rheumatoid arthritis. Nat Med 26, 1295–1306 (2020).

68. Zhang B, et al. CD127 imprints functional heterogeneity to diversify monocyte responses in inflammatory diseases. J Exp Med 219, (2022).

69. Zhang F, et al. IFN-γ and TNF-α drive a CXCL10+ CCL2+ macrophage phenotype expanded in severe COVID-19 lungs and inflammatory diseases with tissue inflammation. Genome Med 13, 64 (2021).

70. Kwon G, Park Y, Kang K, Park-Min KH, Kang K. Epigenomic landscapes define differential Janus kinases inhibitor sensitivity in IFN-gamma-primed human macrophages. iScience 28, 112502 (2025).

