## Supplemental Figures1-7 for "TACE reprograms RANKL-mediated differentiation of macrophages by activating the non-canonical pathway of IRF3"

### Online Supplemental Materials

#### TACE-mediated inhibition of the crosstalk between IRF3 and HBEGF enhances osteoclastogenesis and arthritic bone erosion.

Se Hwan Mun, Brian Oh, Shunichi Yokota, Akio Umemoto, Andrew Suh, Kyuho Kang, Woojung Kim, David Oliver, Tania Pannellini, Geunho Kwon, Young Yang, Liang Deng, and Kyung-Hyun Park-Min

- **Supplemental Figures ------------------------------------ 1- 8**
- **Supplemental Table 1-------------------------------------- 9**

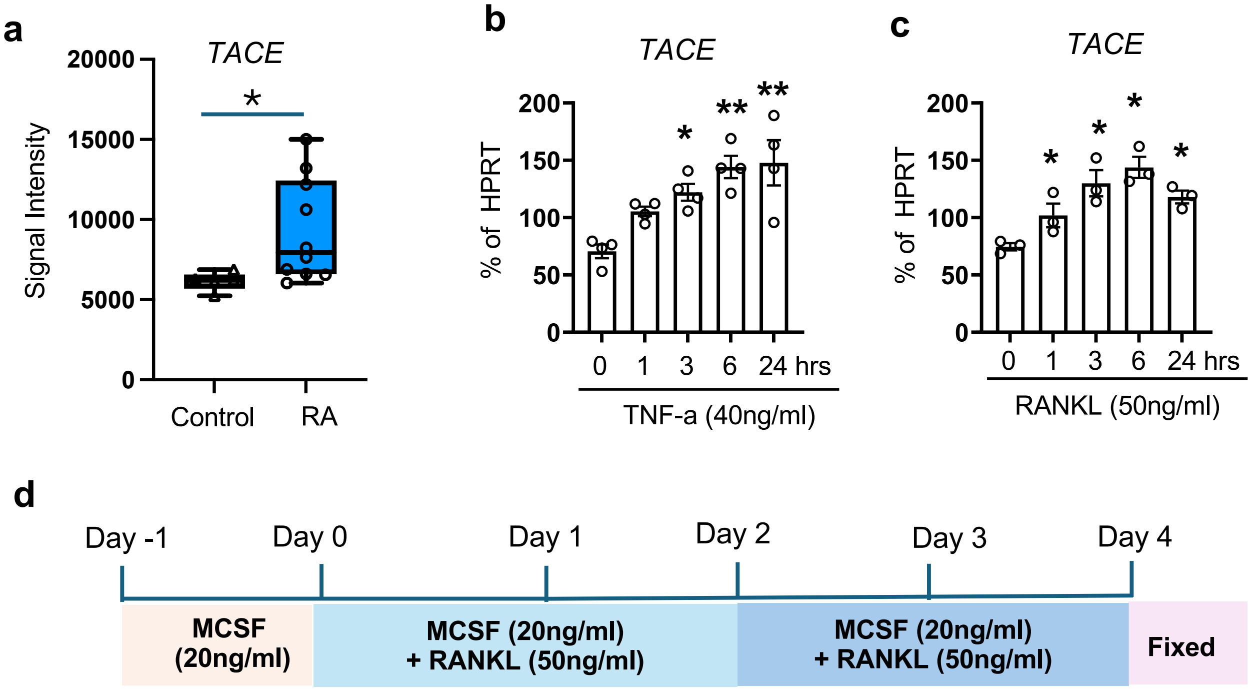


**Supplementary Figure 1. TACE expression is elevated in synovial fluid (SF) CD14⁺ cells from patients with rheumatoid arthritis (RA) and is upregulated by TNF-α and RANK stimulation**. (**a**) Synovial fluid (SF) CD14$^+$ cells were isolated from patients with rheumatoid arthritis (RA, n=9). Control macrophages were cultured with M-CSF for 3 days (n=5). The data were obtained from GSE97779 (**b**, **c**) CD14$^+$ cells isolated from healthy donors were cultured for 24 hours with (**b**) TNF-α (40ng/ml, n=4) or (**c**) RANKL (50ng/ml, n=3). TACE expression levels were measured with RT-qPCR. (d) The schematic illustrating our experimental setup and treatment with M-CSF (20 ng/ml) and RANKL (50 ng/ml) for human osteoclast formation. All data are presented as mean ± SEM. *p < 0.05, **p < 0.01 by One-way ANOVA with *a post hoc Tukey* test (**a** - **c**).

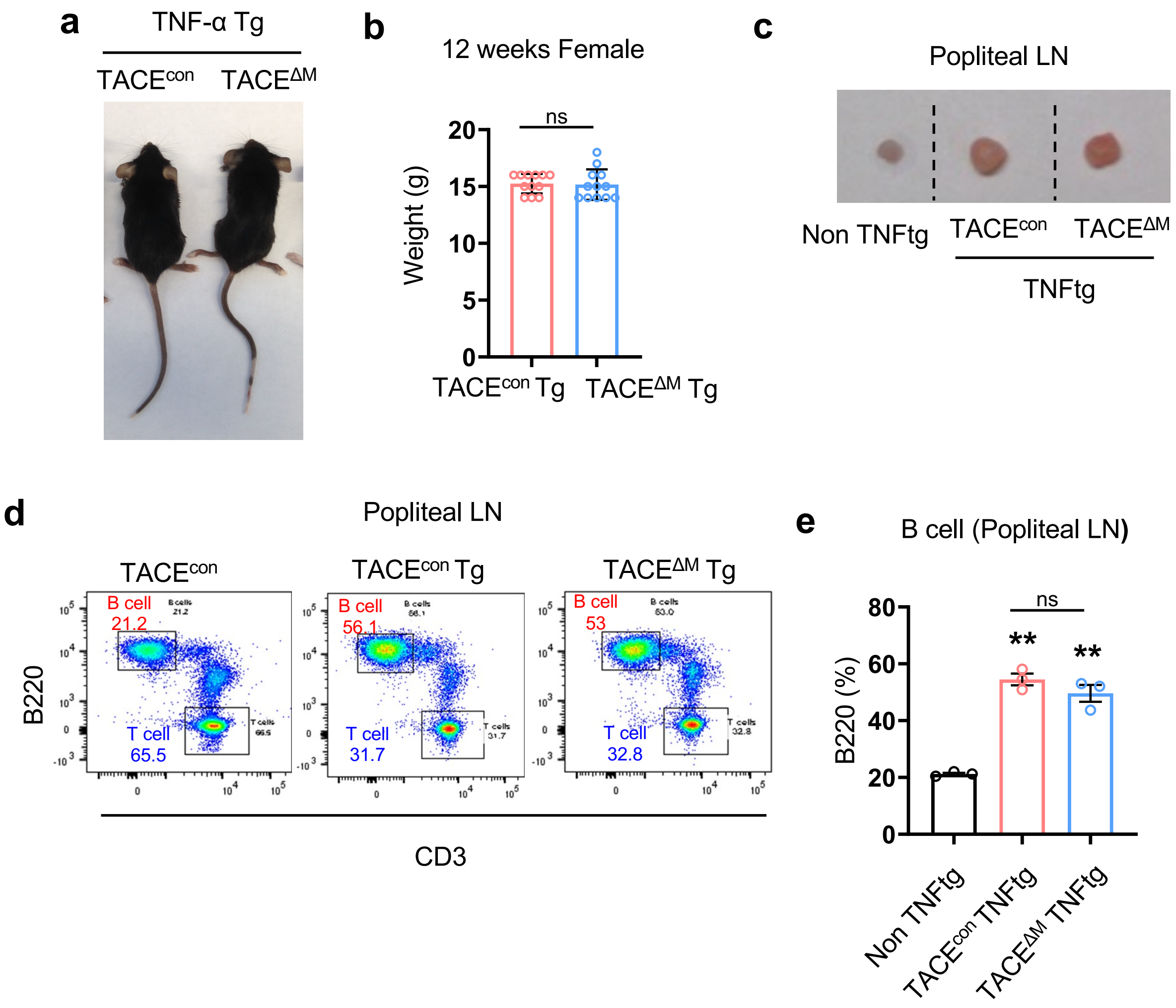


**Supplementary Figure 2. Phenotype of TACE$^{\Delta M}$ TNF-Tg mice and TACE$^{con.}$ TNF-Tg mice**

(**a**) Representative images of 12-weeks-old TACE$^{con}$ TNF-Tg and TACE$^{\Delta M}$ TNF-Tg mice. (**b**) Body weights of TACE$^{con}$Tg and TACE$^{\Delta M}$ Tg (n=12)mice (**c**-**e**) Popliteal lymph nodes were isolated from non-TNFTg, TACE$^{con}$ Tg and TACE$^{\Delta M}$ Tg(n=3) mice. (**c**) The size (**d**, **e**) T and B cell populations in the popliteal lymph node were analyzed by flow cytometry (FACS). All data are presented as mean ± SEM. **p < 0.01 by unpaired *t-test* (**b**) and One-way ANOVA with *a post hoc Tukey* test (**e**).



Supplemental figure 2

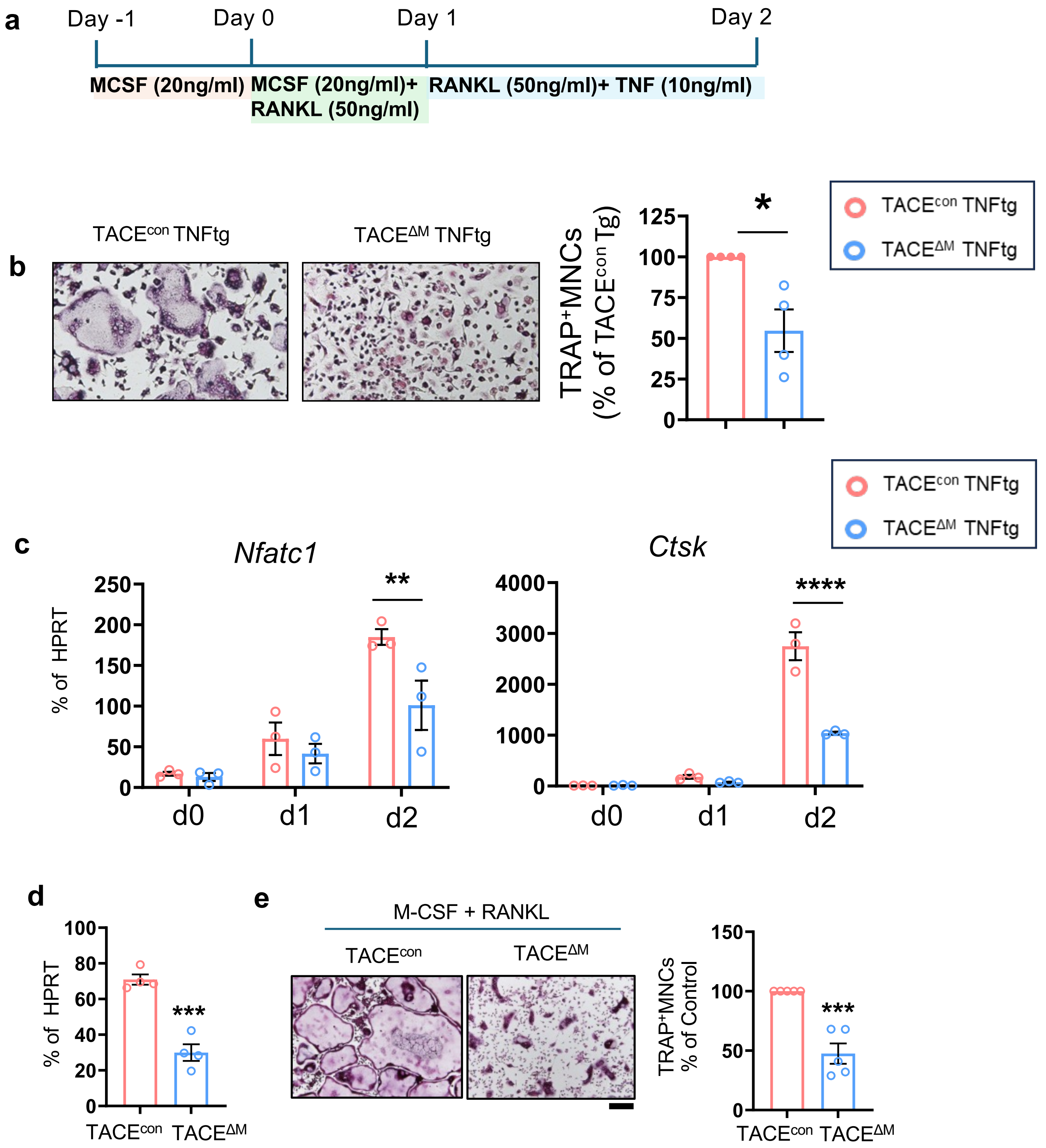




Supplemental figure 3

**Supplementary Figure 3. TACE positively regulates osteoclastogenesis.**

(a-c) Bone marrow-derived macrophage (BMDM) from TACEcon TNF Tg and TACEΔM TNF Tg mice were cultured with M-CSF (20 ng/ml) and RANKL (50 ng/ml). TNF-α (10ng/ml) was added the following day. (a) Experimental schematic for osteoclast formation. (b) Osteoclastogenesis assay. Left: Representative TRAP-stained wells. Scale bar, 100 μm. Right: Quantification of TRAP-positive multinucleated cells (≥3 nuclei), normalized to TACEcon TNF Tg. (c) NFATc1and CSTK mRNA expression in response to RANKL stimulation in TACEcon TNF Tg and TACEΔM TNF Tg cells. (**d**,**e**) Assessment of TACE deletion efficiency and osteoclastogenic capacity in myeloid-specific TACE-deficient cells (TACEfl/fl:LysM Cre mice) under non-TNFtg conditions. (**d**) Confirmation of TACE deletion efficiency, (**e**) Osteoclastogenesis assay. Left: Representative TRAP-stained wells. Scale bar, 100 μm. Right: Quantification of TRAP-positive multinucleated cells (≥3 nuclei), normalized to $TACE^{con}$. (**f**, **g**) Effect of TACE deletion on osteoclastogenesis. Cells derived from the fetal livers of wild-type (WT) and TACE knockout (KO) embryos were cultured with M-CSF and RANKL to induce osteoclast formation. *p < 0.05; **p < 0.01 , ***p < 0.001 by unpaired *t-test*(**b**,**d,e**) and One-way ANOVA with *a post hoc Tukey* test(**c**). Data represent at least three independent experiments.

Supplemental figure 3

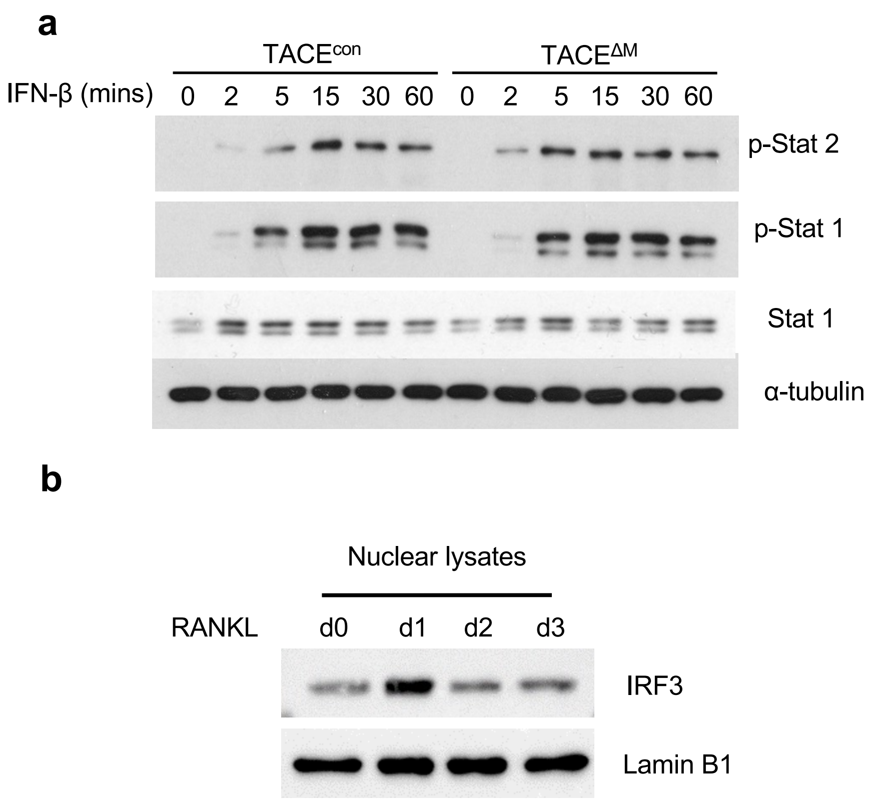


**Supplementary Figure 4. TACE deficiency does not affect the type I IFN signaling.**

(**a**) BMDMs from TACE$^{con}$ and TACE$^{\Delta M}$ mice were stimulated with IFN-β(100 units/ml). Immunoblot analysis using antibodies against phosphorylated STAT1 (p-STAT1), STAT2 (p-STAT2), STAT1 or α-tubulin. (**b**) BMDMs from WT mice were stimulated with RANKL (50 ng/ml) for the indicated times. Nuclear proteins were extracted and analyzed by immunoblotting with IRF3 or Lamin B1 antibodies. Data represent at least three independent experiments.

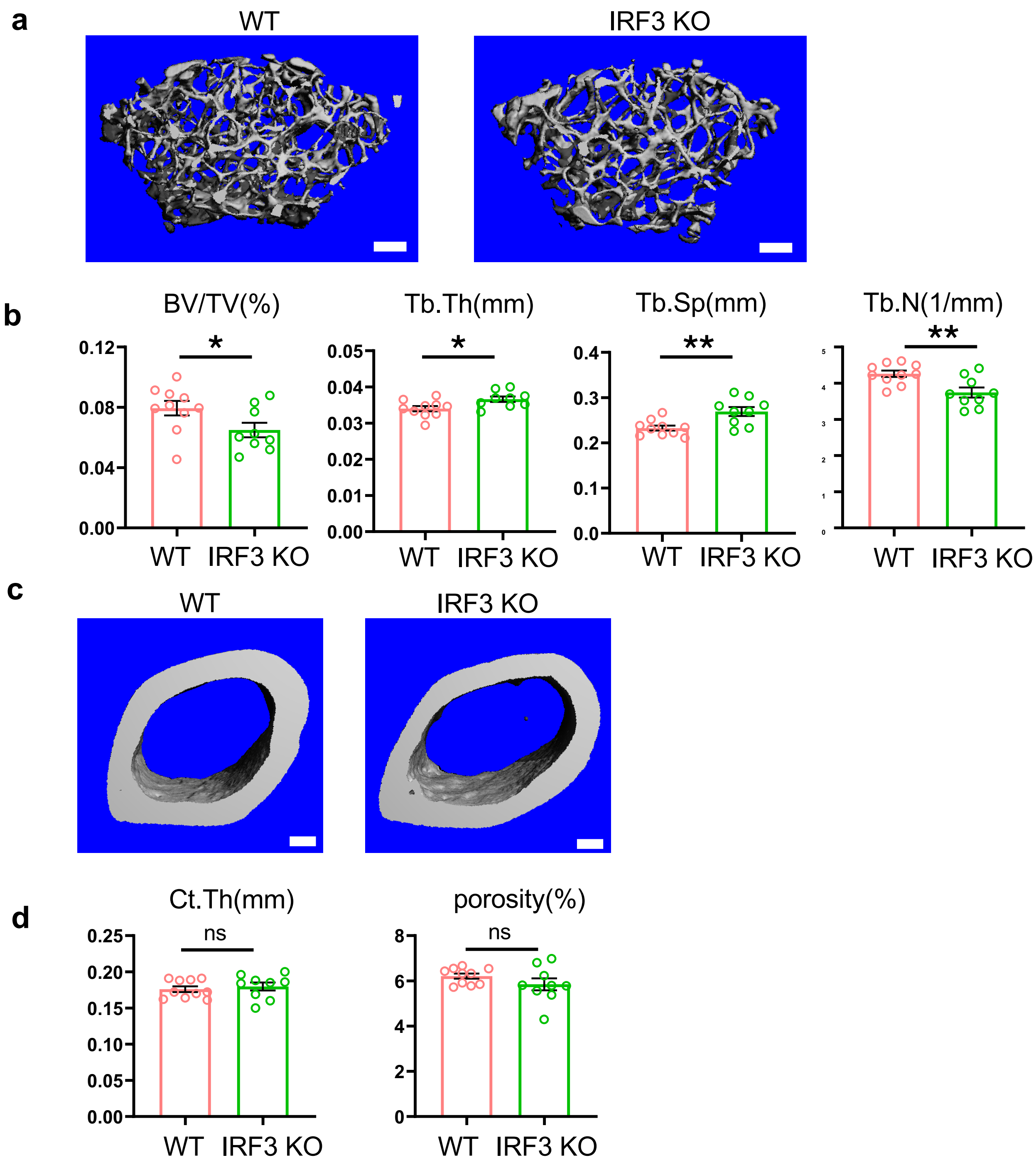


**Supplementary Figure 5. IRF3 is a negative regulator of osteoclastogenesis**

(**a** and **b**) Micro-CT analysis of femurs from 12-week-old WT (n=10) and IRF3KO mice (n = 9). Scale bar: 200μm. (**a**) Representative images of the distal trabecular bone in femurs.(**b**) Analysis of bone parameters in the distal trabecular region. Bone volume/tissue volume ratio (BV/TV), trabecular thickness (Tb.Th), trabecular number (Tb.N), and trabecular separation (Tb.Sp) were quantified by micro-CT.(**c** and **d**) Representative images of the distal cortical bone in femurs. (**d**) Analysis of bone parameters in the distal cortical region. cotical thickness (Ct.Th), porosity. All data are presented as mean ± SEM. $*p < 0.05$; $**p < 0.01$ by unpaired *t-test*(**b,d,f**) Data represent at least three independent experiments

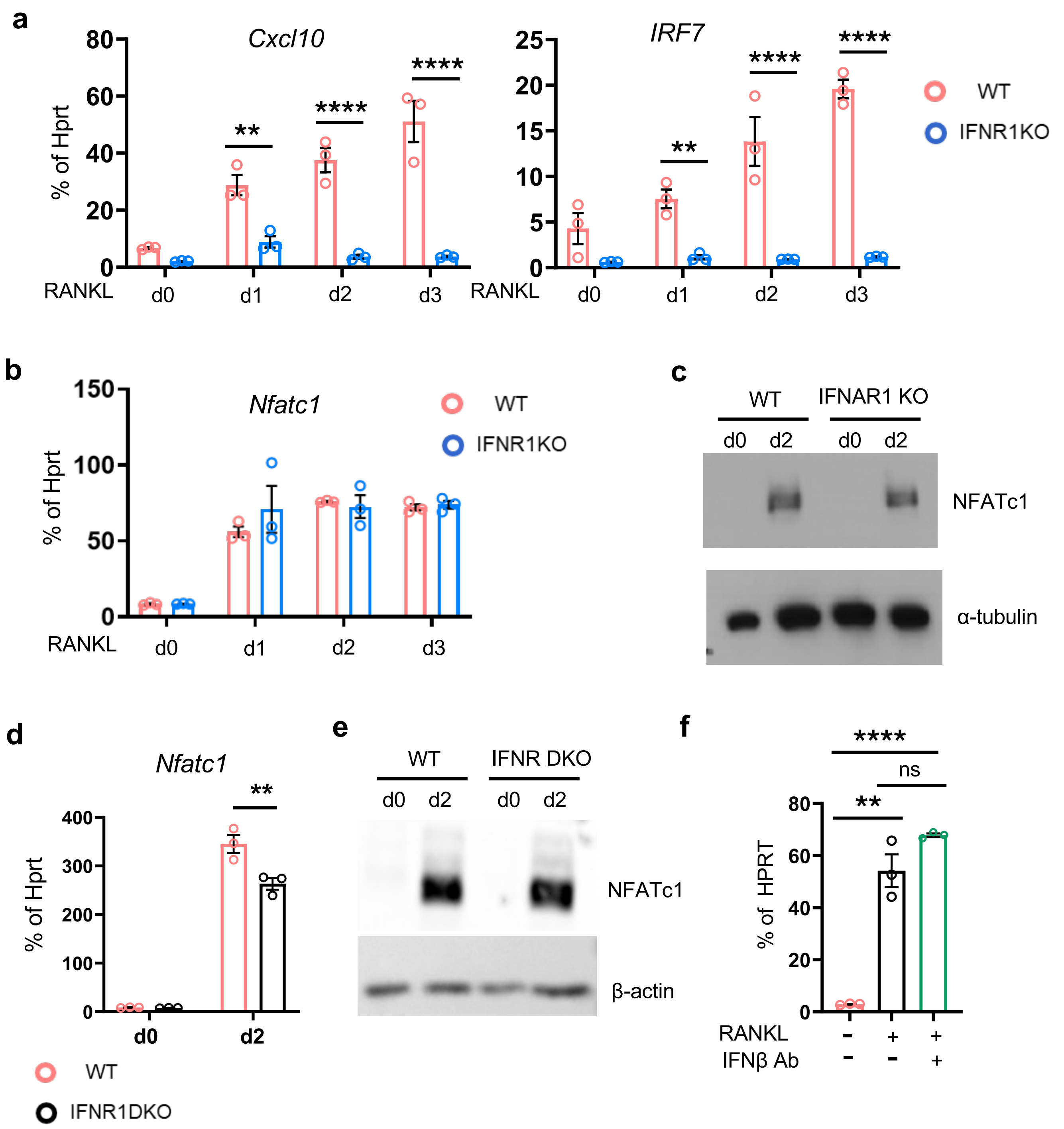


**Supplementary Figure 6. IFNs do not suppress RANKL-induced NFATc1 expression.**

(**a**-**c**) BMDMs from wild-type (WT) and IFNAR1 knockout (KO) mice were cultured with M-CSF and RANKL for 3 days.(**a**) Expression of interferon-stimulated genes (ISGs) in WT and IFNAR1KO cells.(**b**) NFATc1 mRNA expression in response to RANKL stimulation in WT and IFNAR1KO cells.(**c**) Immunoblot analysis using antibodies against NFATc1 and α-tubulin.(**d**, **e**) BMDMs from WT and IFNR DKO (IFNAR1/IFNGR1 double-deficient mice) were cultured with RANKL for 2 days. Both mRNA (**d**) and protein (**e**) of NFATc1 expression. (**f**) WT BMDMs were treated with anti-IFNβ antibodies, and NFATc1 expression was measured by RT-qPCR. All data are presented as mean ± SEM. n.s., not significant. *p < 0.05; **p < 0.01, ***p < 0.005, ****p < 0.001 by one-way ANOVA with *post hoc Tukey's* test (**a**,**b**,**d**,**f**). Data represent at least three independent experiments.

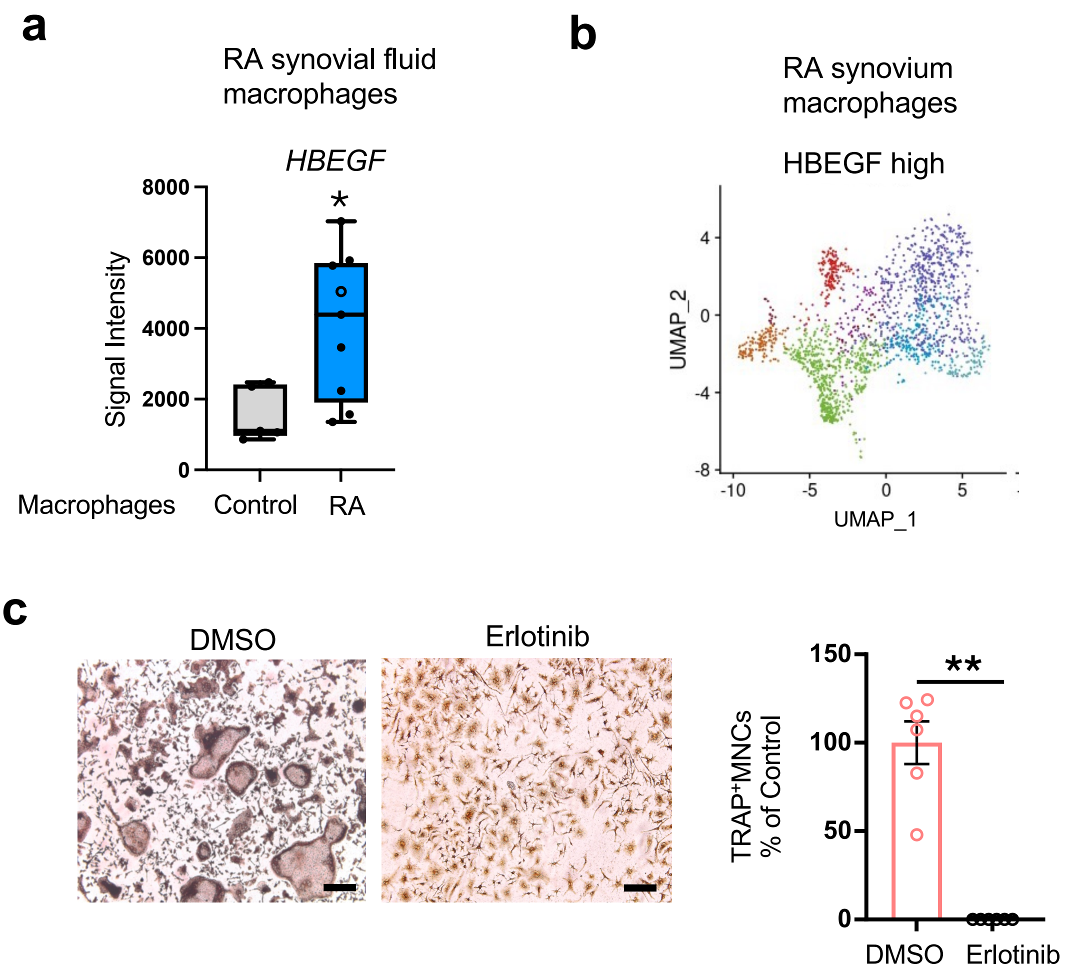


**Supplementary Figure 7. HB-EGF Is Enriched in RA Synovial Macrophages**

**(a, b)** *HBEGF*-high cells are present in synovial fluid macrophages(**a**) [66] and across multiple macrophage subpopulations of RA synovium, though at varying frequencies and expression intensities (**b**). (**b**) The pre-processed gene-cell matrix of the scRNA-seq data [13] was obtained from EMBL's European Bioinformatics Institute (EMBL-EBI) database (Array Express: E-MTAB-8322) and reanalyzed following the methods of a previous study [16] . *HBEGF*-high cells were defined as those with normalized expression values greater than 0.5, and *HBEGF*-low cells as those with expression values equal to or below 0.5. (**c**) BMDMs from WT mice were cultured with M-CSF and RANKL, with or without erlotinib, for 3 days. Osteoclastogenesis assay. *Left*: Representative TRAP-stained wells. Scale bar, 100 μm. *Right*: Quantification of TRAP-positive multinucleated cells (≥3 nuclei), normalized to the number of WT osteoclasts. All data are presented as mean ± SEM. n.s., not significant. *$p < 0.05$; **$p < 0.01$ by unpaired t-test(**a**,**c**). Data represent at least three independent experiments.

.

**Supplementary Table 1**

A list of primers used in this study:

| Gene Symbol | Sequence |
|---|---|
| hNFATc1 | Forward: 5'-CTTCTTCCAGTATTCCACCTAT-3'<br>Reverse: 5'-TTGCCCTAATTACCTGTTGAAG-3' |
| hITGB3 | Forward: 5'-CCCCAAATCCCTCCCCACAAATAC-3'<br>Reverse: 5'-GGAAGAACGCGCCAGAGCAAAATG-3' |
| hCTSK | Forward: 5'-CTCTTCCATTTCTTCCACGAT-3'<br>Reverse: 5'-ACACCAACTCCCTTCCAAAG-3' |
| hTACE | Forward: 5'-ACCCTTTCCTGCGCCCCAGA-3'<br>Reverse: 5'-GTTTTGGAGCTGCTGGCGCC-3' |
| hIRF3 | Forward: 5'-ACACATACTGGGCAGTGAGC-3'<br>Reverse: 5'-CTACAATGAAGGGCCCCAGG-3' |
| hGAPDH | Forward: 5'- ATCAAGAAGGTGGTGAAGCA-3'<br>Reverse: 5'-GTCGCTGTTGAAGTCAGAGGA-3' |
| hHPRT | Forward: 5'-GACCAGTCAACAGGGGACAT-3'<br>Reverse: 5'-CCTGACCAAGGAAAGCAAAG-3' |
| mNfatc1 | Forward: 5′- CCCGTCACATTCTGGTCCAT-3′<br>Reverse: 5′- CAAGTAACCGTGTAGCTCCACAA -3′ |
| mItgb3 | Forward: 5'-CCGGGGGACTTAATGAGACCACTT-3'<br>Reverse :5'-ACG CCC CAA ATC CCA CCC ATA CA-3' |
| mCtsk | Forward: 5′- CGTGGGAGACATGACCAGTG-3′<br>Reverse: 5′- CGAACCACACTGGCCCTGGT-3′ |
| mTace | Forward: 5′- CCCCCACCCGGAGATGCTGA-3′<br>Reverse: 5′- CGGCACACACGGGCCAGAAA-3′ |
| mCxcl10 | Forward: 5′- CCAAGTGCTGCCGTCATTTTC-3′<br>Reverse: 5′- GGCTCGCAGGGATGATTTCAA-3′ |
| mIrf7 | Forward: 5′- GGGACCTCTTGCTTCAGGTT-3′<br>Reverse: 5'- AAACACGGTCTTGCTCCTGG-3' |
| mIrf3 | Forward: 5′- GGGATCCTGAACCTCGTTCG-3′<br>Reverse: 5'-CAATTCCTCCCCTGGCTAGA-3' |
| mMx1 | Forward: 5′-GGCAGACACCACATACAACC-3′<br>Reverse: 5′-CCTCAGGCTAGATGGCAAG-3′ |
| mHbegf | Forward: 5′- AGGACCTGAGCTATAGGAACC-3′<br>Reverse: 5′- CCCAGTCAGGGTAGCAACTG-3′ |
| mHprt | Forward: 5'-TCCTCAGACCGCTTTTTGCC-3'<br>Reverse: 5'-CTAATCACGACGCTGGGACT-3' |
